# A UVB-responsive common variant at chr7p21.1 confers tanning response and melanoma risk via regulation of the aryl hydrocarbon receptor gene (*AHR*)

**DOI:** 10.1101/2021.03.25.436649

**Authors:** Mai Xu, Lindsey Mehl, Tongwu Zhang, Rohit Thakur, Hayley Sowards, Timothy Myers, Lea Jessop, Alessandra Chesi, Matthew E Johnson, Andrew D Wells, Helen T Michael, Patricia Bunda, Kristine Jones, Herbert Higson, Rebecca C Hennessey, Ashley Jermusyk, Michael A Kovacs, Maria Teresa Landi, Mark M Iles, Alisa M Goldstein, Melanoma Meta-Analysis Consortium, Jiyeon Choi, Stephen J Chanock, Struan F A Grant, Raj Chari, Glenn Merlino, Matthew H Law, Kevin M Brown

## Abstract

Genome-wide association studies have identified a melanoma-associated locus on chromosome band 7p21.1 with rs117132860 as the lead SNP, and a secondary independent signal marked by rs73069846. rs117132860 is also associated with tanning ability and cutaneous squamous cell carcinoma (cSCC). As ultraviolet radiation (UVR) is a key environmental exposure for all three traits, we investigated the mechanisms by which this locus contributes to melanoma risk, focusing on cellular response to UVR. Fine-mapping of melanoma GWAS identified four independent sets of candidate causal variants. A GWAS region-focused Capture-C study of primary melanocytes identified physical interactions between two causal sets and the promoter of the aryl hydrocarbon receptor gene (*AHR*). Subsequent chromatin state annotation, eQTL, and luciferase assays identified rs117132860 as a functional variant and reinforced *AHR* as a likely causal gene. As AHR plays critical roles in cellular response to dioxin and UVR, we explored links between this SNP and *AHR* expression after both 2,3,7,8-tetrachlorodibenzo-p-dioxin (TCDD) and ultraviolet B (UVB) exposure. Allele-specific AHR binding to rs117132860-G was enhanced following both, consistent with predicted weakened AHR binding to the risk/poor-tanning rs117132860-A allele, and allele-preferential *AHR* expression driven from the protective rs117132860-G allele was observed following UVB exposure. Small deletions surrounding rs117132860 via CRISPR abrogates AHR binding, reduces melanocyte cell growth, and prolongs growth arrest following UVB exposure. These data suggest *AHR* is a melanoma susceptibility gene at the 7p21.1 risk locus, and rs117132860 is a functional variant within a UVB-responsive element, leading to allelic *AHR* expression, and altering melanocyte growth phenotypes upon exposure.

## Introduction

Cutaneous melanoma (CM) is the deadliest form of skin cancer^1^, and it arises from melanocytes, pigment producing cells in the skin. Ultraviolet radiation (UVR) exposure is a well-established environmental risk factor^2^, while genetics plays a clear role, with twin studies suggesting melanoma to be the most heritable of solid tumors^3^. Over the past decade, a series of progressively larger genome-wide association studies (GWAS) have identified 68 independent signals at 54 genomic loci to be associated with cutaneous melanoma at a level of genome-wide significance^4–14^, highlighting the role of genetics in disease risk. Of these loci, many have also been implicated in risk-associated complex traits, including nevus count^8,14–16^ and multiple pigmentation phenotypes ^14,17–21^. Notably, an analysis by Landi and colleagues found that of these 54 loci, almost half (n=26) were also significantly associated with ease of tanning in UK Biobank participants ^14^.

Among these tanning- and melanoma-associated loci is a risk locus on chromosome band 7p21.1 between the *AGR3* and *AHR* genes, initially identified by Law and colleagues in a meta-analysis of GWAS data from Europe, Australia and United States^4^. The association to this region has since been confirmed by a larger meta-analysis, which identified two independent genome-wide significant signals, as well as a third following conditional analyses ^14^, all of which lie closest to *AGR3*. The strongest association signal at this locus has been found to be the lead signal reported to be associated with both tanning response to sun exposure14,22 as well as cutaneous squamous cell carcinoma (cSCC), where the risk alleles for melanoma and cSCC were found to be associated with an inability to tan. Thus, these data suggest that response to ultraviolet radiation is the common underlying genetic etiology for these traits.

Amongst the potential candidate causal genes at this locus is the gene encoding the aryl hydrocarbon receptor (*AHR*), a ligand-activated transcription factor expressed in all skin cell types ^24,25^. It is well characterized as a pleiotropic sensor of environmental factors in response to UVR, dioxins, polycyclic aromatic hydrocarbon (PAH), tryptophan and its metabolites, and cigarette smoke ^25–28^. Upon activation, AHR translocates to the nucleus and forms a complex with Aryl Hydrocarbon Receptor Nuclear Translocator (ARNT), which binds to DNA recognition sequence and initiates gene transcription ^25,29^. Notably, AHR signaling has been reported to modulate melanogenesis in mice via multiple mechanisms, inducing hyperpigmentation in response to benzanthrone^30^, mediating UVB-induced skin tanning in mice via changing expression of genes involved in melanocyte proliferation and differentiation including *kitl*^26^, or regulating melanoma-stroma interaction ^31^. AHR has also been found to play a key role in melanoma cell behavior, where *AHR* expression was associated with MEK inhibitor efficacy in *NRAS*-mutant cell lines^32^. Sustained activation of AHR signaling by exogenous and endogenous signals, including UVR, leads to a transcription signature which is similar to that of metastatic and dedifferentiated melanoma cells, as well as that of BRAF inhibitor resistance. Chronic activation of the canonical AHR signaling pathway can switch BRAF inhibitor sensitive cells into persister/resistant cells, emphasizing the importance of AHR antagonists as potential sensitizers for melanoma target therapy. [32]

In this report, we establish the regulation of the *AHR* gene by the melanoma susceptibility locus at 7p21.1 marked by SNP rs117132860. In melanocytes, we provide evidence for a chromatin interaction between the region surrounding rs117132860 and the promoter and gene body of *AHR gene*, as well as similar interactions from candidate causal variants from a second independent signal at this locus. Chromatin immunoprecipitation (ChIP) by AHR indicated allele-specific binding to the protective rs117132860-G allele, which was enhanced in response to UVB and 2,3,7,8-tetrachlorodibenzo-p-dioxin (TCDD) exposure, accompanied by an increase of *AHR* expression. CRISPR knock out of rs117132860 abolished the AHR binding to this region and notably alters melanocyte proliferation and cellular response to UVB exposure. In summary, the association of chr7p21.1 with both melanoma risk and tanning response implies a mechanistic link between two processes, and we show this occurs through AHR, providing a direct link between germline genetics and environmental factors on cancer risk.

## Results

### Fine-mapping identifies rs117132860 as the lone candidate causal SNP representing the primary melanoma association signal at 7p21.1

Landi and colleagues ^14^ recently reported the results from a meta-analysis of melanoma GWAS for 36,760 cases, providing evidence that multiple common sequence variants on chromosome band 7p21.1 confer melanoma susceptibility. The most significant association was for rs117132860 (*P*_meta_ = 3.83 x 10^-21^, OR_G-allele_ = 0.71), with an apparent second signal ~138kb away marked by rs73069846 (*P*_meta_ = 1.24 x 10^-8^, OR_C-allele_ = 0.95; linkage-disequilibrium [LD] r^2^_rs117132860_ = 0.0004, D’_rs117132860_ = *0.13*, 1000G EUR; **Fig. 1, Supplementary Fig. 1**); this second signal appears to represent that locus/SNP previously identified by Law and colleagues in a smaller melanoma meta-analysis (rs1636744; r^2^_rs73069846_ = 0.83; D’_rs73069846_ = 0.94; r^2^_rs117132860_ = 0.0003; D’_rs117132860_ = 0.11; 1000G EUR)^4^. Additionally, conditional- and joint-analysis identified a third independent signal of risk-associated variants marked by rs10487582 (*P*_conditional_ = 2.30 x 10^-8^). Notably, rs117132860 had previously been reported to be the lead signal in this region associated with other UVR-associated traits, including tanning ability (*P* = 7.63 x 10^-23^; OR_G-allele_ = 1.30) ^22^ as well as cutaneous squamous cell carcinoma (*P* = 3.6 × 10^-8^; OR_G-allele_ = 0.68) ^23^. Subsequent analyses by Landi and colleagues of nevus count^15^, as well pigment trait data from UK Biobank^14^ found significant or marginal associations for both the primary and secondary melanoma signals (nevus count: *P*_rs117132860_ = 6.53 x 10^-7^, *P*_rs73069846_ = 0.00277; UKBB tanning: *P*_rs117132860_ = 6.60 x 10^-91^, *P*_rs73069846_ = 1.92 x 10^-21^; UKBB skin color: *P*_rs117132860_ = 3.64 x 10^-45^, *P*_rs73069846_ = 3.86 x 10^-15^; UKBB childhood sunburns: *P*_rs117132860_ = 2.97 x 10^-9^, *P*_rs73069846_ = 0.00261) ^14^, suggesting a direct link at this locus between UV exposure, melanoma, and these melanoma-associated traits.

**Fig. 1:**
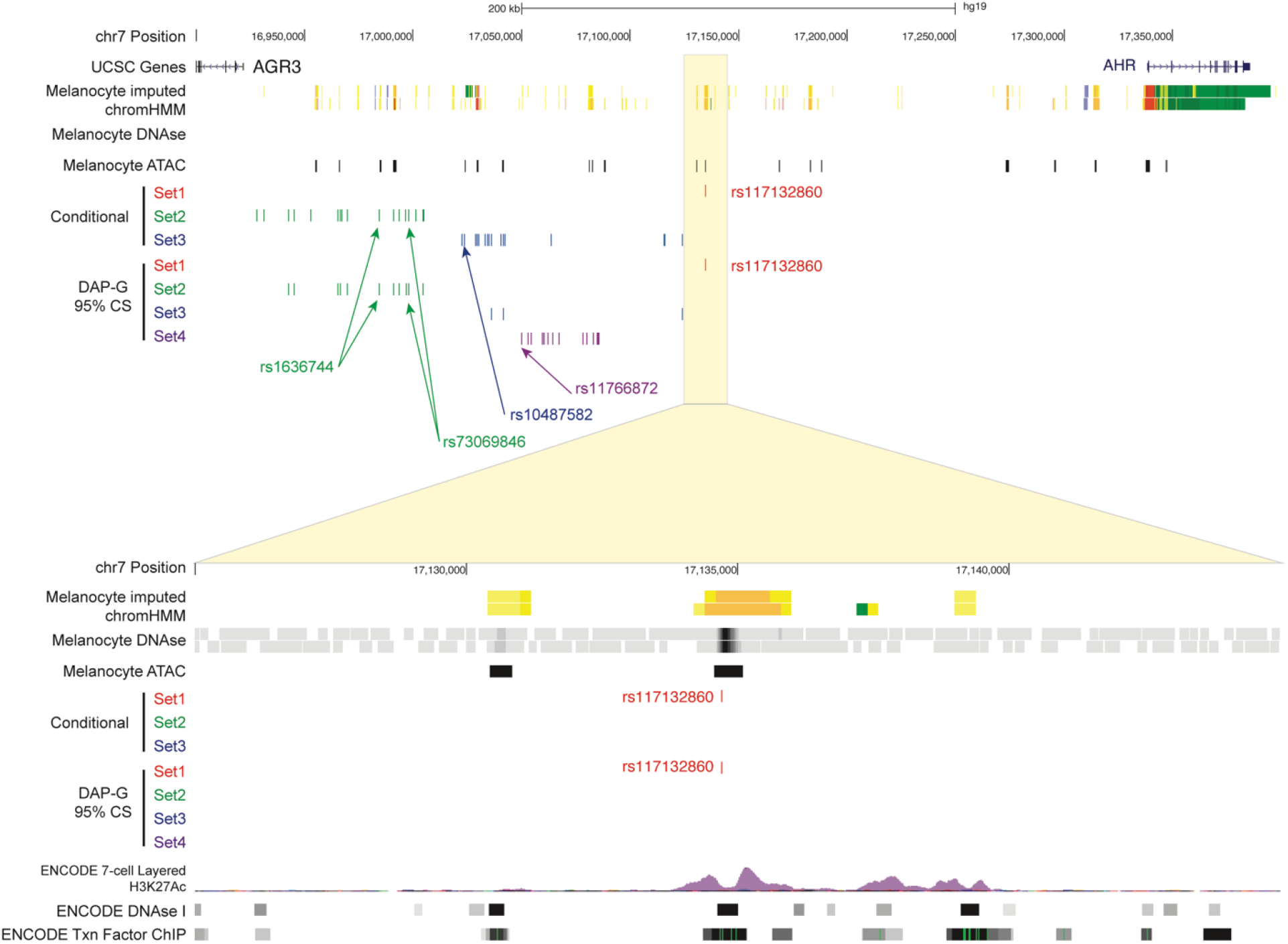
Fine mapping of multiple melanoma GWAS signals on chromosome band 7p21.1. (Top) a view of the 7p21.1 locus including UCSC genes, imputed ChromHMM and DNaseI hypersensitivity (DHS) data from two melanocyte cultures generated by the RoadMap Project, ATAC-seq data generated from five human primary melanocyte cultures, and candidate causal SNP sets nominated by either conditional analysis or Bayesian fine-mapping (95% credible sets for each of four clusters) using DAP-G. Candidate causal variants mapping to signal/cluster 1 are in red, signal/cluster 2 in green, signal/cluster 3 in blue, and signal/cluster 4 in purple. (Bottom) a zoomed-in view of the region immediately surrounding rs117132860, showing rs117132860 mapping to open chromatin and annotated melanocyte enhancer. Genomic positions are based on hg19. For the zoomed-in region containing rs117132860 (bottom), layered 7-cell H3K27Ac, DHS clusters, and transcription factor CHIP tracks were generated by ENCODE and provided through the UCSC Genome Browser.

To identify credible causal melanoma risk variants within this larger region, we performed both conditional association analyses, as well as Bayesian fine-mapping. Firstly, we performed conditional analysis iteratively for each of the three previously identified association signals using a leave-one-out approach, conditioning on the lead SNP of the remaining two signals for each in order to obtain independent credible sets of variants representing each signal. The signal marked by rs117132860 had no other SNP within two orders of magnitude log likelihood relative to the lead *P*-value following conditional analysis (rs117132860 *P*_conditional_ = 2.99 x 10^-20^; conditioned on rs73069846 and rs10487582; **Supplementary Table 1, Fig. 1, Supplementary Fig. 1, Supplementary Fig. 2a**), suggesting rs117132860 as the lone candidate causal variant for this signal. The secondary signal consisted of a set of 17 variants within two orders of magnitude log-likelihood ratio relative to the lead SNP (rs73069846 *P*_conditional_ = 1.44 x 10^-10^; conditioned on rs117132860 and rs10487582; **Supplementary Table *2*, Fig. 1, Supplementary Fig. 1, Supplementary Fig. 2b**), while the third set consisted of 16 such variants relative to the lead (led by rs9638738, *P*_conditional_ = 1.10 x 10^-8^; conditioned on rs117132860 and rs73069846; **Supplementary Table 3, Fig. 1, Supplementary Fig. 1, Supplementary Fig. 2c**). Lastly, conditioning on all three resulted in a marginal signal marked by rs11766872 (*P*_conditional_ = 4.81 x 10^-6^; **Supplementary Table 4; Supplementary Fig. 2d**). We also performed Bayesian fine mapping of the 7p21.1 region (+/- 250 kb centered on rs117132860) using DAP-G ^33,34^, allowing for up to four causal variants given the results of the conditional analysis. Here, the analysis identified four clusters of candidate causal variants (**Fig. 1, Supplementary Fig. 1, Supplementary Tables 5-6**). The first consisted solely of rs117132860 (Set 1; PIP = 1), while a second cluster consisted of a 95% credible set of 11 variants (Set 2; marked by rs73069846, PIP = 0.25). The third 95% credible set consisted of three variants marked by rs34585474 (Set 3; PIP = 0.84), with the fourth cluster consisting of a set of 14 SNPs (Set 4; led by rs12670784, PIP = 0.32). Finally, we assessed whether there may be other credible causal variants not successfully imputed in the meta-analysis or assessed in conditional or Bayesian analyses. Notably, the meta-analysis by Landi and colleagues performed imputation using a Haplotype Reference Consortium (HRC) reference panel that lacked small insertion/deletion variants. Thus, we also sought to identify additional variants not reported in the meta-analysis but in high LD with the lead SNP for each signal using LDLink (https://ldlink.nci.nih.gov/)^35^. We identified no high-LD proxy variants (r^2^>0.6; 1000G EUR) for rs117132860, two indel variants in LD to rs73069846 (Signal 2; rs35785866 and rs200020478), none for rs34585474, and five for rs12670784 (Signal 4; rs13229759, rs199662382, rs10532327, rs367629062, rs10589929) for (1000G EUR) that were not already imputed in our dataset. These data collectively suggest that there are multiple association signals within this locus, map to four sets of potential credible causal variants, and indicate that rs117132860 is the sole candidate causal variant for the signal most strongly associated with melanoma.

Functional annotation using human primary melanocyte ATAC-seq data we generated from a set of five genetically-independent cultures, as well as additional melanocyte histone-ChIPseq and DNaseI hypersensitivity sequencing data from the Roadmap Epigenomics Project^36^ and ENCODE ^37^, indicated that rs117132860 lies in an open chromatin region marked as an active enhancer in primary cultured skin melanocytes (**Fig. 1, Supplementary Fig. 1, Supplementary Table 7**). Furthermore annotation of candidate causal SNPs in the region for predicted alteration of transcription factor binding using motifbreakR^38^ indicated that rs117132860 is within an AHR::ARNT binding motif with the risk (A) allele weakening predicted binding (**Fig. 2a, Supplementary Table 8**). Notably, the gene encoding AHR itself lies 200 kb away from rs117132860 (**Fig. 1, Supplementary Fig. 1**), suggesting a potential role for feedback regulation of AHR levels through rs117132860. Candidate causals for other signals also overlap marks indicating potential melanocyte *cis*-regulatory regions, including rs847404 (ATAC-open), rs975603 (ATAC-open) and rs847377 (DNaseI open) for DAP-G Set 2, rs847428 (ATAC-open) and rs847429 (ATAC-open) for DAP-G Set 3, and rs4721562 (ATAC-open) for DAP-G Set 4 (**Supplementary Table 9**; Set 1 highlighted in pink, Set 2 in yellow, Set 3 in green, Set 4 in blue). Taken together, these data strongly suggest that rs117132860 is the lone candidate causal variant representing the primary melanoma risk association signal at chr7p21.1, suggest that rs117132860 may also represent a potential causal variant for other melanoma-associated traits, and nominate several strong candidate causal variants for independent melanoma GWAS signals in the region.

**Fig. 2:**
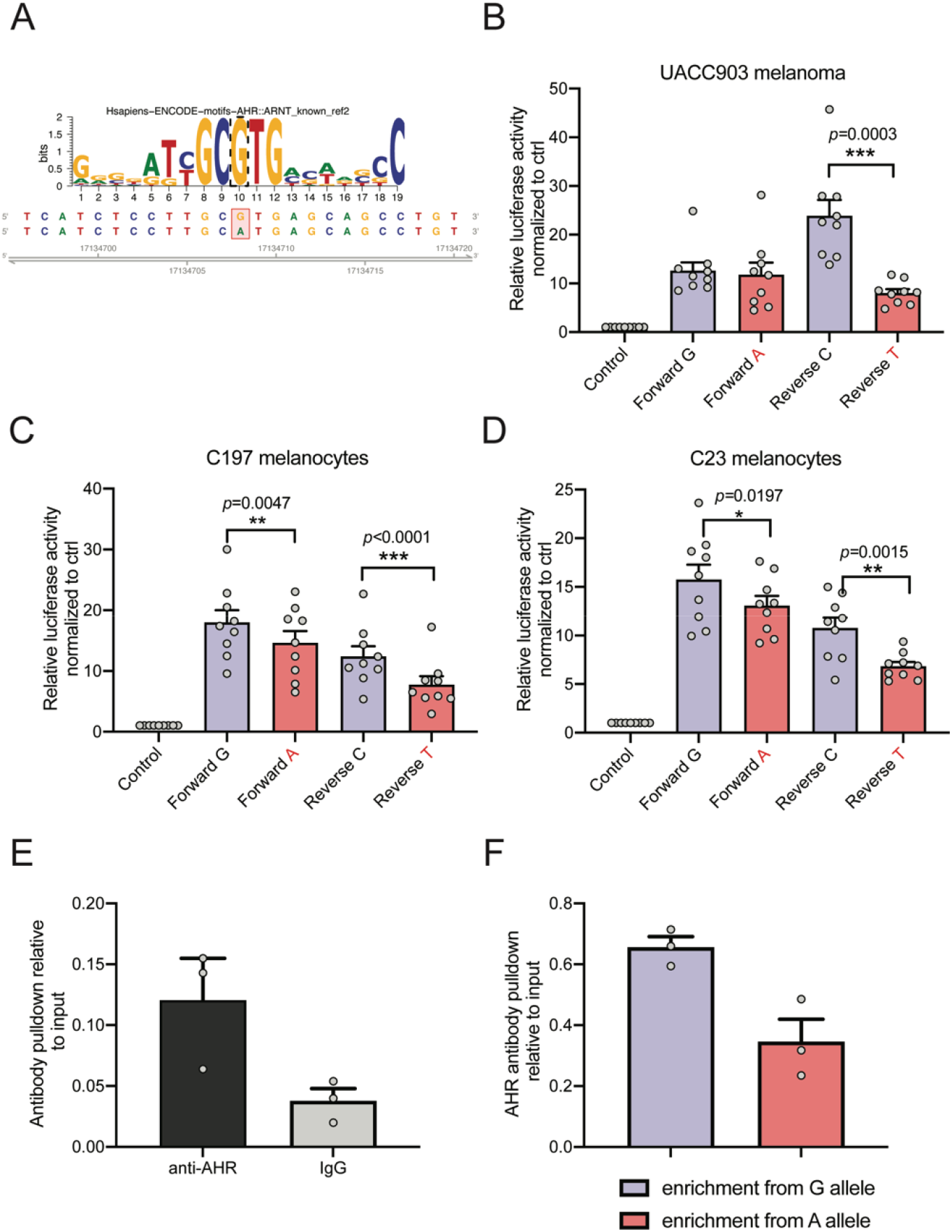
Allele-specific transcriptional regulation and AHR binding via rs117132860. (A) motifbreakR analysis identified a AHR::ARNT binding motif as strongly altered, where the melanoma-protective rs117132860-G allele matches the motif, while rs117132860-A is predicted to alter AHR binging. (B-D) Luciferase reporter activity of a 375bp fragment encompassing either allele of rs117132860 is measured in (B) the human melanoma cell line UACC903 and (C,D) two primary melanocyte cultures (C197 and C23; labeled). pGL4.23 is the control vector containing a minimal TATA promoter; risk (labeled in red) and protective alleles of rs117132860 are assayed in both forward and reverse directions relative to the promoter. Three biological replicates were combined for each cell (three technical replicates of each, mean, and SEM are plotted); (E) rs117132860 displays allele-preferential binding to AHR in C87, melanocytes (heterozygous for rs117132860) by chromatin immunoprecipitation assay using anti-AHR antibody or normal IgG control. Relative quantities are shown as fold over input DNA (one representative experiment from three biological replicates is shown); (F) Taqman genotyping of rs117132860 using AHR ChIP DNA from the same experiment was normalized to input DNA (one representative set from 3 biological replicates is shown; mean of PCR triplicates with SEM is plotted).

We also assessed whether any of these candidate causal variants demonstrated allele-specific reporter activity in a previously published massively parallel reporter assay (MPRA) analysis of the region ^39^. As these MPRA data assessed allelic activity for SNPs in LD with the lead SNP (rs1636744) from the prior meta-analysis by Law and colleagues (2015)^4^, the SNPs tested largely corresponded to Set 2, while rs117132860 was not assessed as this signal had not yet been found to be genome-wide significant. Four candidate causals, however, were FDR-significant (FDR < 0.01) in a joint analysis of UACC903 and HEK cells, including two variants where the risk allele exhibited higher reporter activity (rs73069846 and rs847404) and two where the risk allele showed lower expression (rs847377 and rs9648229; **Supplementary Table 9**). Two of these also overlap melanocyte regulatory marks: rs847404 (open chromatin in melanocyte ATAC-seq); and rs847377 (DNaseI accessible in melanocytes from RoadMap Epigenome Project).

### rs117132860 is located within a melanocyte enhancer and displays allele-specific *cis*-regulatory activity

As described above, functional annotations indicate that rs117132860 is localized to an active melanocyte enhancer region (**Fig. 1**), and itself alters an AHR binding motif (**Fig. 2a**) and is the sole candidate causal variant for the lead melanoma association signal. Thus, to assess the gene regulatory potential of this region as well as allelic differences in *cis*-regulatory activity, we cloned a 375bp fragment encompassing the melanocyte DNaseI hypersensitivity peak region harboring rs117132860 into a luciferase reporter plasmid and assessed luciferase activity in melanoma cells as well as human cultured primary melanocytes. Compared to control plasmid lacking this region, the fragment containing the genomic region surrounding rs117132860 displays strong transcriptional activity in both forward and reverse directions in UACC 903 melanoma cells (**Fig. 2b**) and two independent melanocyte cultures (C197 and C23) (**Fig. 2c-d**), consistent with this region acting as a transcriptional enhancer. Notably, in human primary melanocytes, we observed that the risk-associated A allele of rs117132860 exhibited significantly lower reporter activity than the protective G allele when cloned in both forward and reverse direction (**Fig. 2c-d**; C197 *P* = 0.0047 and *P* = 8.09 x 10^-6^, C23, *P* = 0.0197 and *P* = 0.0015; F and R respectively; two-tailed Student’s T-test). In melanoma cells, this allele-specific reporter activity was significant only when the fragment was cloned in reverse direction (**Fig. 2b**, *P* = 0.0003; two-tailed Student’s T-test). Collectively, these data are consistent with the region surrounding rs117132860 acting as an enhancer in cells of melanocytic lineage and suggest that rs117132860 alters the *cis*-regulatory activity of this enhancer in an allele-specific manner, with lower expression associated with the risk allele.

### rs117132860 genotype is correlated with expression of *AHR*

We next assessed expression quantitative trait locus data (eQTL), both to assess rs117132860 as a potential *cis*-regulatory variant, as well as to potentially link this variant to regulation of a candidate causal gene(s) within the topologically-associated domain (TAD) harboring rs117132860. We turned to an eQTL dataset we previously generated from cultured primary human melanocytes (n=106)^40^, where a genome-wide significant eQTL was not previously observed for rs117132860. Given low power due to the small sample size of this dataset and the relatively low minor allele frequency for rs117132860 (MAF = 0.017, 1000G EUR), we assessed whether rs117132860 might nonetheless be an eQTL for any of the five genes within the TAD. Of these, *AGR2* and *AGR3* were poorly expressed (detectable expression in <20% of melanocyte cultures, RSEM >0; **Fig. 3a**). Of the remaining, we found a strong correlation between the A-risk allele and lower levels of *AHR* expression (*P* = 0.00136, slope = 0.715; **Fig. 3b**). The remaining genes exhibited weak correlations which were not significant after multiple-testing correction for the number of genes tested within the TAD (**Fig. 3c-d**; *TSPAN13, P* = 0.03; *BZW2, P* = 0.10). Notably, the direction of correlation for *AHR* is consistent with reporter assay data where higher expression levels were associated with the protective rs117132860-G allele, providing further support that rs117132860 may be *cis*-regulatory and linking the risk allele to lower levels of *AHR*. Finally, while rs117132860 is not a genome-wide significant eQTL in any tissues assessed by the Genotype and Tissue Expression Project (GTEx), assessment of predicted *AHR* levels modeled from GTEx data via transcriptome-wide association study (TWAS) by the TWAS-Hub (http://twas-hub.org/genes/AHR/) show a correlation between modeled *AHR* levels and tanning ability^41^, further suggesting a role for common *AHR*-regulatory sequence variants and tanning.

**Fig. 3:**
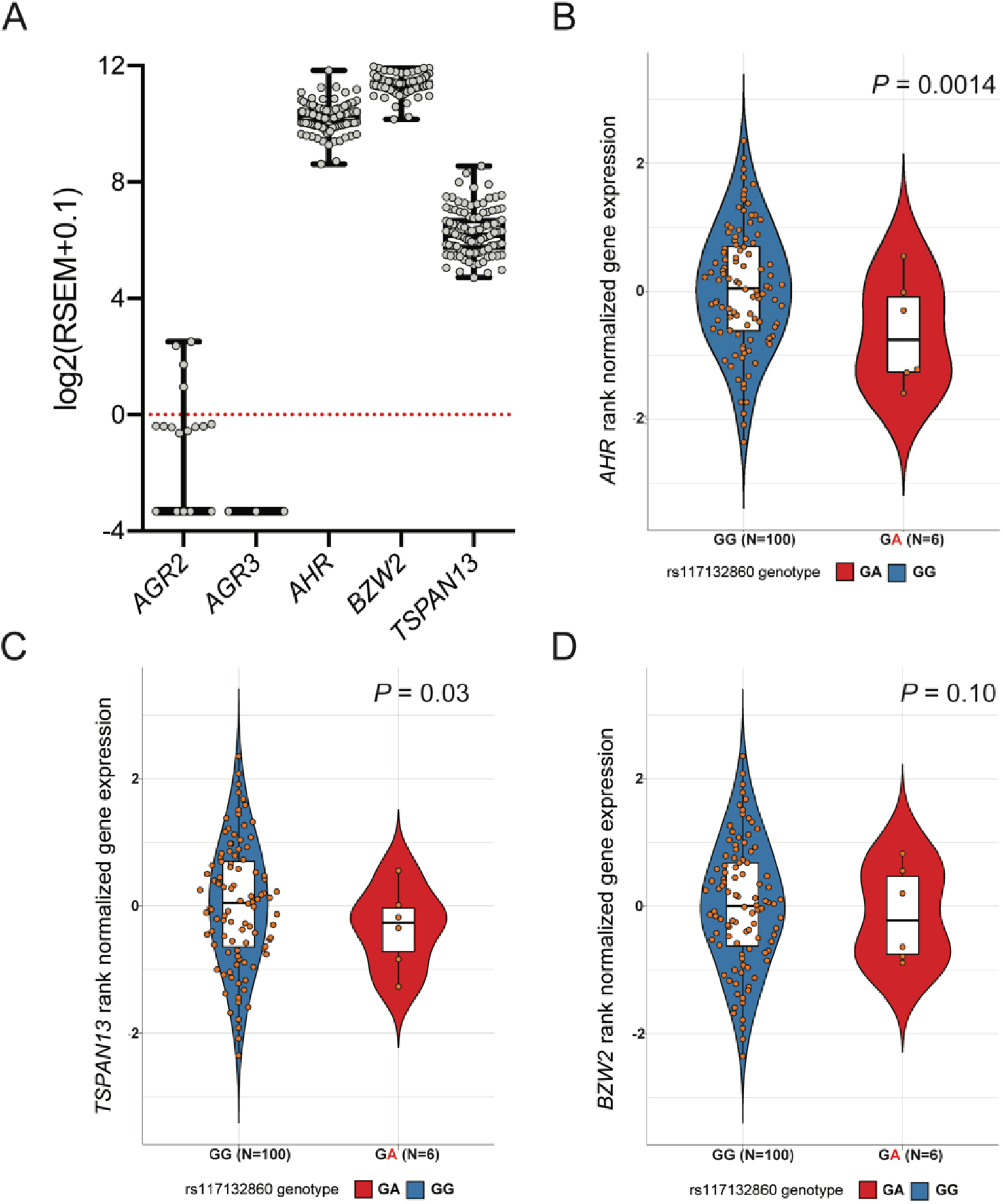
rs117132860 genotype is associated with *AHR* expression in human melanocytes. (A) Expression of the five genes located in the TAD containing rs117132860 in 106 human primary melanocyte cultures as assessed by RNA-sequencing (box and whiskers with Min to Max). (B-D) Rank-normalized gene expression of (B) *AHR*, (C) *TSPAN13* and (D) *BZW2* in melanocytes with GG or GA genotype at rs117132860. For violin plots, center lines show the medians; box limits indicate the 25th and 75th percentiles; whiskers extend 1.5 times the interquartile range from the 25th and 75th percentiles, all samples are represented by dots. n = 100 (GG), 6 (GA) sample points.

### The enhancer harboring rs117132860 physically interacts with the *AHR* promoter and gene body

We next assessed physical interactions between regions of association and nearby genes using multiple methods. As a part of a larger, region-specific Capture-C assay, we designed baits targeting the regions of association for the two signals at chromosome band 7p21.1 that were genome-wide significant in single-SNP analyses from the most recent meta-analysis by Landi and colleagues based on single-SNP data (set 1 and set 2; **Fig. 4a; Supplementary Fig. 3; Supplementary Table 10**)^14^. Using these baits, we performed Capture-C assays for a set of five genetically distinct primary human melanocyte cultures (3 biological replicates per culture). Following loop calling by CHiCAGO^42^ using all replicates of all five melanocyte cultures grouped together, we assessed the targets of chromatin looping between our regions of association (and fine-mapped SNPs) to genes within the TAD harboring rs117132860. Notably, we observed a direct physical contact between the restriction fragment containing rs117132860 and melanocyte-promoter (annotated by melanocyte imputed chromHMM data) region for *AHR* (**Fig. 4a; Supplementary Fig. 3; Supplementary Tables 9, 12 and 13**), suggesting that the enhancer containing rs117132860 may regulate AHR. We did not observe physical association between rs117132860 and any other gene within the TAD containing rs117132860, suggesting *AHR* is the primary target of this enhancer. We subsequently confirmed the physical association between rs117132860 and AHR using chromatin conformation capture (3C), using the region harboring rs117132860 as probe and designing primers spanning from rs117132860 3’ to past the *AHR* gene body. 3C analysis in three independent melanocyte cultures confirmed a physical association between rs117132860 and the *AHR* promoter, as well as the *AHR* gene body (**Fig. 4b**), further implicating *AHR* as a target gene of the enhancer containing rs117132860.

**Fig. 4:**
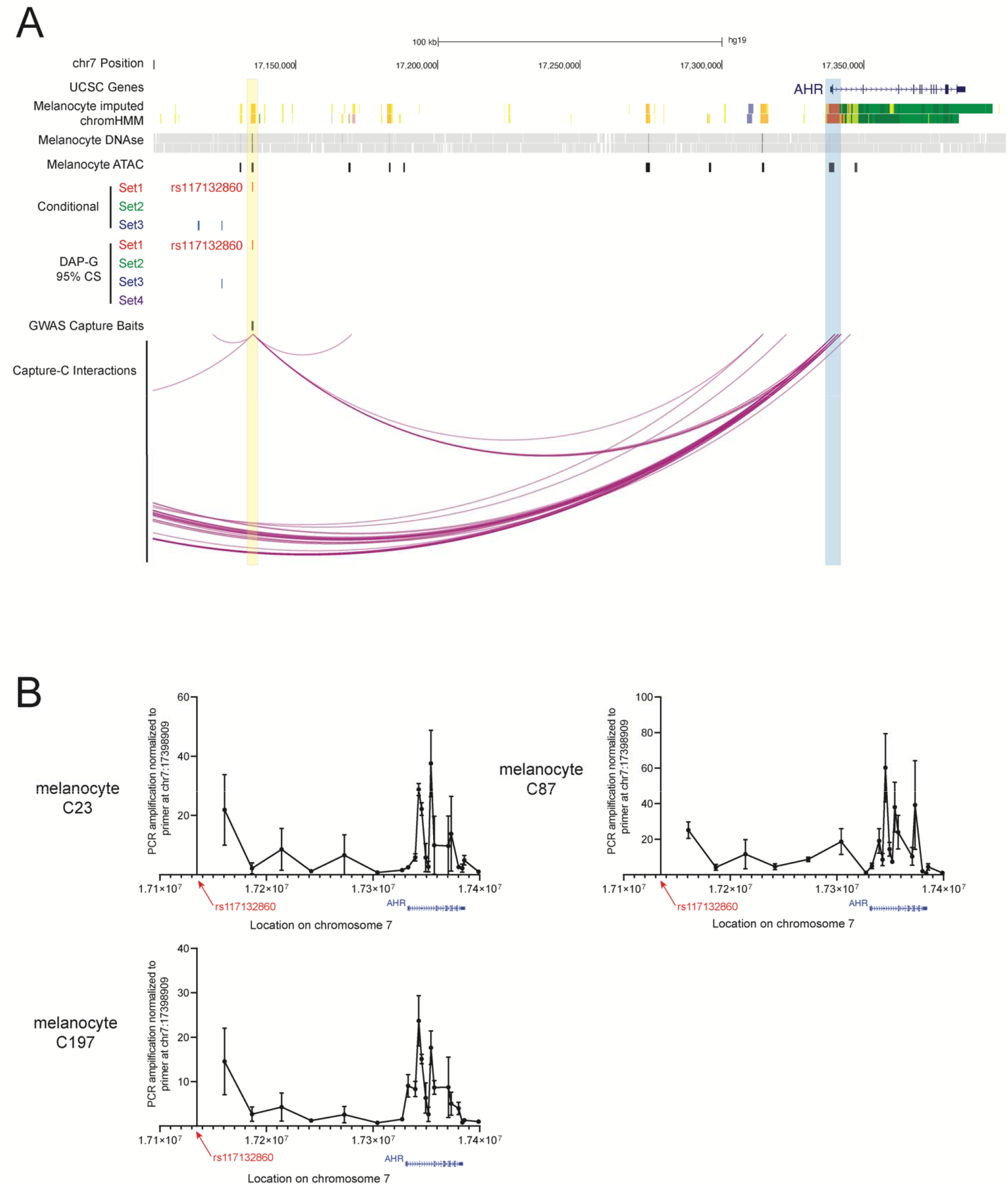
Region-specific capture-C and chromatin conformation capture (3C) show a chromatin interaction between rs117132860 and the AHR promoter and gene body. (A) Significant chromatin interactions captured by Capture-C between rs117132860 and *AHR* gene. Region capture baits are labeled in black and significant interactions are shown as purple arcs. UCSC genes, imputed ChromHMM and DNaseI hypersensitivity (DHS) data from two melanocyte cultures generated by the RoadMap Project, ATAC-seq data generated from five human primary melanocyte cultures, and candidate causal SNP sets nominated by either conditional analysis or Bayesian fine-mapping (95% credible sets for each of four clusters) using DAP-G are also shown. Loops were called from data from five distinct melanocyte cultures (3 biological replicates per culture) analyzed together in order to detect the most reproducible interactions. (B) Interactions between rs117132860 and *AHR* were confirmed via chromatin conformation capture (3C) in three independent primary melanocyte cultures. Relative interaction frequencies of various genomic fragments to the rs117132860 region are shown according to their location in chromosome 7. For each experiment, the PCR amplification for each target primer from 3C libraries were first normalized to the PCR amplification of the target primer from BAC library DNA, and subsequently normalized to the PCR amplification of the target primer HindIII17398909, which is immediately downstream of the *AHR* gene body. A total of four biological replicates were performed (one each for C23 and C87, two for C197).

In addition to the chromatin loop between the *AHR* gene and rs117132860, we observed multiple chromatin loops between the secondary conditional/DAP-G signal at this locus and the promoter of *AHR* (**Supplementary Fig. 3; Supplementary Table 9, 12-13**), suggesting that candidate causal variants from multiple independent association signals in this region target *AHR*. Specifically, we see direct loops between fragments containing DAP-G Set 2 SNPs rs847404, rs1721040, rs12535242, and rs35785866 which was not fine mapped but is in LD with rs73069846. Taken together, these data show a physical association between multiple association signals within the 7p21.1 melanoma risk locus and the *AHR* promoter, strongly implicating *AHR* as the likely causal gene explaining the melanoma risk association for this larger region.

### Dynamic change of *AHR* expression and allele-specific binding of AHR to rs117132860-G in melanocytes upon dioxin and UVB exposure

As described previously, motif analysis indicates that rs117132860 is part of the AHR binding motif ‘GCGTG”, with the risk-A allele weakening the binding relative to the protective-G allele (**Fig. 2a**). AHR footprinting using data from multiple cell types including epidermal melanocytes also identified rs117132860 as within a region likely to be bound by AHR (RegulomeDB; https://regulomedb.org/regulome-search/)^43^. We therefore assessed the potential binding of rs117132860 by AHR via chromatin immunoprecipitation (ChIP) assay in primary melanocyte cultures. As shown in **Fig. 2e**, an AHR antibody specifically pulled down a fragment harboring rs117132860 compared to IgG control as assessed by quantitative real-time PCR (qRT-PCR), suggesting AHR indeed binds to this fragment. A qRT-PCR genotyping assay was further designed using specific probes distinguishing between rs117132860-G and -A alleles. In a melanocyte culture heterozygous for rs117132860, enhanced IP by AHR antibody was detected for the G allele relative to the A allele (**Fig. 2f**), consistent with the motif prediction. Together with the luciferase result, these data indicate that rs117132860 may regulate transcription in an allele specific manner as part of an AHR transcription factor binding site.

AHR is activated in response to a number of environmental factors, including dioxins and UVR, upon which it translocates to the nucleus and forms a complex with aryl hydrocarbon receptor nuclear translocator (ARNT), subsequently binding to the AHR DNA recognition sequence and initiates target gene transcription. Given this mode of activation, along with the relatively weak binding of AHR to rs117132860 observed in normal culturing conditions (**Fig. 2e**), we assessed *AHR* expression and AHR-binding to the enhancer harboring rs117132860 in the context of dioxin (2,3,7,8-Tetrachlorodibenzo-p-dioxin; TCDD) and ultraviolet B (UVB) exposure in multiple human primary melanocyte cultures. We observe that both transcription of *AHR*, as well as overall cellular and nuclear levels of AHR protein are changed dynamically following TCDD and UVB exposure. Following TCDD exposure, *AHR* mRNA expression is induced from 3 to 24 hours post-exposure (**Fig. 5a, Supplementary Fig. 4a**) while we observe an increase in nuclear AHR via Western blot at 1 hour post exposure, followed by a reduction in both whole cell and nuclear AHR from 2 to 6 hours post exposure (**Fig. 5b**); it has previously been reported that whole cell AHR protein level decreases after TCDD treatment in human melanocytes while *AHR* mRNA level increases ^27^. This difference between mRNA and protein level could be mediated by posttranslational ubiquitin-mediated protein degradation, similar to what has been reported following FICZ treatment^44^. Following UVB exposure (13.2 mJ/cm^2^), on the other hand, we observe an increase in the nuclear level of AHR starting at 4 hours and decrease of cytosolic level around 8-24 hours following exposure, accompanied by increases in *AHR* mRNA from 24-72 hours. (**Fig. 5c-d, Supplementary Fig. 4b**). In summary, upon exposure with both TCDD and UVB in melanocytes, AHR protein translocates into the nucleus together with a first decrease then increase in *AHR* expression.

**Fig. 5:**
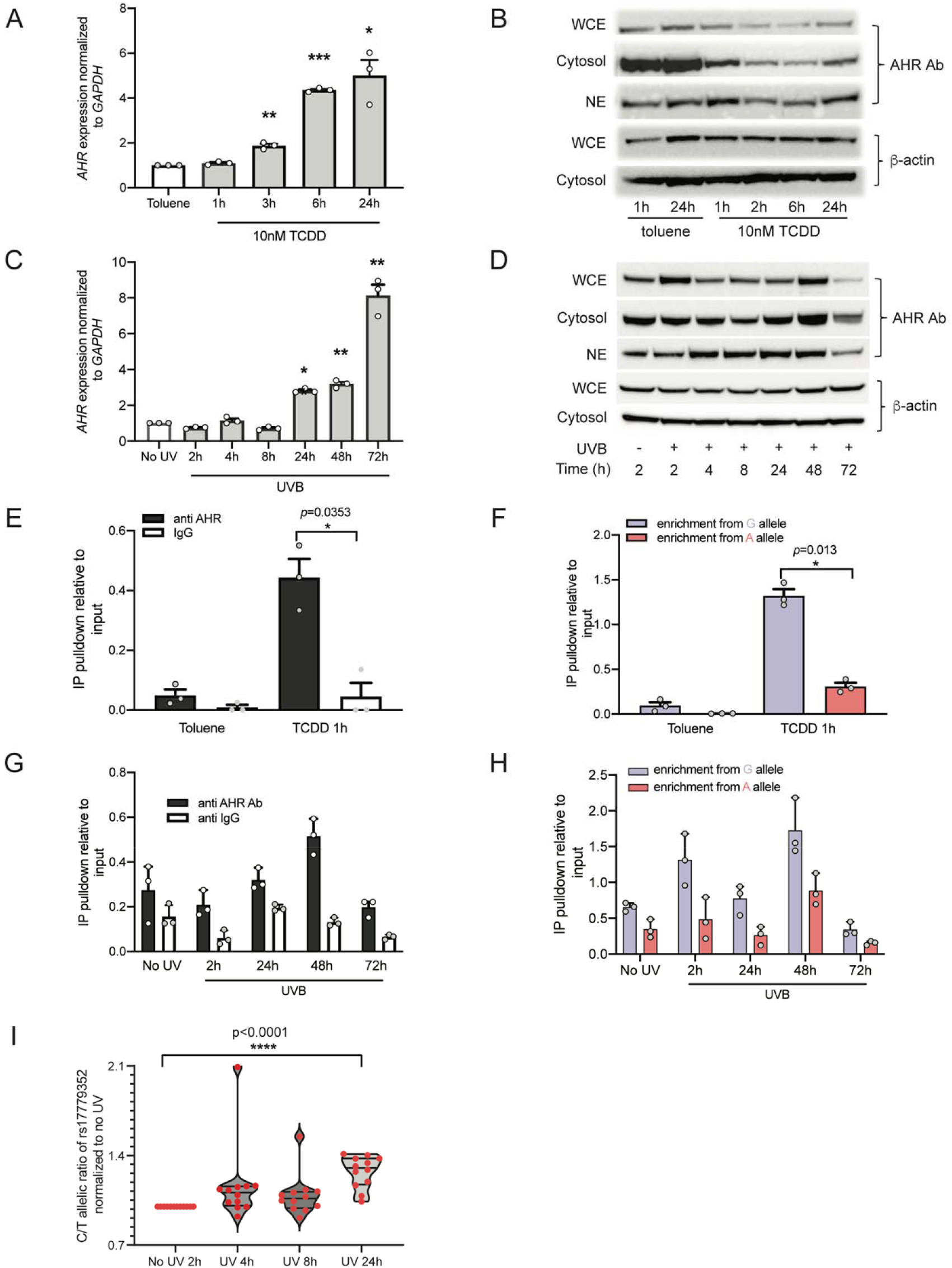
Dynamic change of *AHR* expression and allele-preferential AHR binding to rs117132860-G in melanocytes upon dioxin and UVB exposure. (A) *AHR* expression normalized to *GADPH* was measured by Taqman assay in C87 human melanocytes before and after TCDD treatment. (B) AHR western blotting indicates a dynamic change of AHR protein nuclear localization after TCDD treatment. (C) *GAPDH* normalized *AHR* transcription is increased after UVB exposure in C87 melanocytes. (D) Nuclear translocation of AHR protein is observed after UVB exposure via western blotting in C87 melanocytes. (E, G) AHR binding to rs117132860 measured by CHIP assay both increased after (E) TCDD treatment in C197 melanocytes and (G) UVB exposure in C87 melanocytes. *: *P* < 0.05; **: *P* < 0.01; ***: *P* < 0.001. A genotyping assay using AHR CHIP DNA shows the enhanced binding of AHR to the rs117132860-G allele both after (F) TCDD treatment and (H) UVB exposure in C87 melanocytes. (I) A genotyping assay of rs17779352, a proxy SNP for rs117132860 located in in the *AHR* coding sequence, indicates that the ratio of AHR expression from the melanoma-protective rs117132860-G/rs17779352-C allele significantly increased after UVB exposure relative to the rs117132860-A/rs17779352-T allele at 24 hours. All experiments except for those shown in (I) were done in both C87 and C197 human melanocytes with a total of four biological replicates each. The experiment set shown in the Fig. is a representative one with *P* value calculated from a two-way paired Student’s T-test (A, C, E, and F) or two-way ANOVA (G and H) and the mean with SEM is plotted. The proxy SNP genotyping experiment (I) was only performed in C87 cells which is heterozygous for both rs117132860 and rs17779352; the violin plot shows the combination of 4 experiments of 3 replicates each with all points shown.

Given these changes in *AHR* transcription as well as AHR nuclear localization, we assessed binding of AHR to the region harboring rs117132860 via AHR chromatin immunoprecipitation (ChIP) both with and without TCDD or UVB exposure in human primary melanocytes. Consistent with the data presented above, in the absence of TCDD or UVB treatment, we observe enriched binding of AHR to rs117132860 via enhanced pulldown by AHR antibody compared to control IgG (**Fig. 5e,g, Supplementary Fig. 5a,c**). At one hour post-exposure, we observe significant increased binding of AHR to rs117132860 relative to untreated toluene control, consistent with the short-term changes in nuclear translocation of *AHR* in response to TCDD and increased expression pattern reflected in latter time points (*P* = 0.035 **Fig. 5e**; and *P* = 0.018, **Supplementary Fig. 5a**). We further assessed allele-preferential binding of AHR following TCDD exposure in primary melanocytes heterozygous for rs117132860 via a Taqman-based allele discrimination assay using dye-labelled probes distinguishing between the G or A allele. Consistent with the predicted stronger AHR motif for rs117132860-G allele, we observe enhanced binding of AHR to rs117132860-G versus rs117132860-A both with and without TCDD treatment, where the preferential binding to G allele becomes significant after TCDD treatment (*P* = 0.013, **Fig. 5f**; and *P* = 0.0014, **Supplementary Fig. 5b**; two-way paired Student’s t-test). Likewise, following UVB exposure, we observe stronger binding of AHR to rs117132860, with the peak binding detected at 48 hours post-exposure (**Fig. 5g, Supplementary Fig. 5c**) and binding of AHR at all timepoints is enriched for the rs117132860-G allele relative to rs117132860-A (**Fig. 5h, Supplementary Fig. 5d**), with both significant after UVB exposure for two melanocyte cultures (*P* = 0.0005, *P* = 2.62 x 10^-6^, respectively for **Fig. 5h**; *P* = 1.3 x 10^-5^, *P* = 7.2 x 10^-12^ for **Supplementary Fig. 5d**; two-way ANOVA). Thus, the dynamic change of AHR binding to the SNP are consistent with the nuclear localization of AHR following both TCDD and UVB exposure and increased expression pattern reflected at later time points, suggesting that activated AHR binding to the SNP by those treatments may regulate *AHR* expression in an allelic manner.

Given the widespread effects of TCDD and in particular UVB on global gene expression, including many control genes, which may affect the quantification of transcription, we turned to assessing allelic expression in human primary melanocyte cultures heterozygous for rs117132860. Specifically, we identified an mRNA coding proxy SNP in LD with rs117132860 (rs17779352; *r^2^ = 0.0014*; D’ = 1, 1000G EUR), which allowed us to assess transcription of *AHR* in *cis* by rs117132860-A and rs117132860-G alleles. We treated a primary melanocyte culture heterozygous for both rs117132860 and rs17779352 (where the A-risk allele of rs117132860 is *cis* to rs17779352-T-allele, and rs117132860-G is *cis* to rs17779352-C) with either TCDD or UVB (13.2 mJ/cm^2^) and collected RNA from cells at different timepoints. Notably, at 24 hours following UVB treatment, we observed a significant increase in *AHR* transcribed from the protective (rs117132860-G, rs17779352-C) allele (P =1.1 x 10^-5^; two-way paired Student’s t-test) compared to the risk allele (**Fig. 5i**). However, we did not observe the same allele-preferential changes in *AHR* expression in response to TCDD (**Supplementary Fig. 6**), These data are consistent in direction with both the luciferase reporter data and *AHR* eQTL with rs117132860. Thus, in response to UVB exposure, we observe an increase in transcription of *AHR* driven from the melanoma-protective rs117132860-G allele as compared to that driven from the rs117132860-A risk allele.

### A melanocyte gene-based CRISPR knockout screen identifies AHR as a gene hit mediating cell growth in immortalized human melanocyte culture

We performed a gene-based pooled CRISPR/Cas9 cell proliferation knockout screen in an immortalized human melanocyte culture, C283T. Here, we designed 3,052 guide RNAs (gRNAs; **Supplementary Table 14**) covering a set of 288 genes, as well as ~200 non-targeting gRNAs. The 288 genes focused primarily on gene candidates from genome-wide significant melanoma GWAS loci. Both Cas9 and gRNAs were expressed in lentiviral vectors, and Cas9 was induced by doxycycline (dox) treatment. DNA samples were collected from sgRNA infected cells before dox treatment (around day 7 after infection) and at three different timepoints after that (around day 14, day 21, and day 28 following infection). gRNAs were sequenced and normalized to counts of the integrated gRNA sequence from cells collected prior to dox treatment. We compared dox-treated cells from all three latter timepoints and prioritized gene hits critical for cell growth and or survival using MAGeCK^45^. By combining three replicate multi-timepoint experiments and comparing read counts (**Supplementary Table 15**) from all samples treated with dox with untreated samples, 55 genes were ranked by MAGeCK as negatively selected genes and two genes as positively selected with significance score <0.05 and FDR<0.05 (**Supplementary Table 16**). Consistent with established roles as tumor suppressors, guides targeting both *TP53* and *CDKN2A* were found to be positively selected, while those targeting *PARP1* and *SETDB1*, two genes previously reported to be important in melanogenesis ^46,47^, are among the top negatively selected gene group. Amongst the negatively selected genes, *AHR* was ranked eighth (log_2_(fold change) = −1.2515), thus suggesting that the *AHR* gene plays a role in melanocyte growth and that risk-associated SNPs at the 7p21.1 play a role in mediating melanocyte cellular phenotypes.

### Deletion of the AHR binding motif encompassing rs117132860 alters melanocyte response to UVB exposure

Finally, in order to further assess the contributions of the enhancer harboring rs117132860 on both expression of *AHR*, and melanocyte cellular phenotypes, we utilized CRISPR-Cas9 editing to mutate/delete the conserved AHR-binding motif containing rs117132860. We designed a sgRNA targeted to rs117132860. A Cas9-expressing lentiviral vector pCW-Cas9-Blast was virally introduced into an TERT-immortalized human melanocyte culture, C283-T, followed by further introduction of either a sgRNA targeted to rs117132860 or a non-target sgRNA. Following selection, we isolated monoclonal cell lines from the mixed population through limited dilution. For candidate clones grown from single cells, we sequenced the genomic region around rs117132860 and deconvoluted the sequences to identify potential cell clones with deletion/mutation regarding to AHR-binding motif around rs117132860. We identified two clones with deletion/mutation of both alleles of the AHR-binding motif around rs117132860 (KO; deletion size in two knockout clones ranged from 2 to 9 bp), one clone with one copy of wild type sequence and one copy with deletion of AHR binding motif (HT; deletion of 16 bp), and 3 clones with no change of the sequence around the SNP (WT) (**Fig. 6a, Supplementary Fig. 7**). We first checked AHR binding to the SNP region in KO cells, and as expected, AHR does not bind to the deleted rs117132860 region both before and after UVB treatment (**Supplementary Fig. 8**). We assessed *AHR* expression from both WT and KO clones with and without UVB exposure. While we see a similar dynamic change of *AHR* expression pattern in WT and KO cells in response to UVB as observed in primary melanocytes, we did not observe a consistent difference in *AHR* expression between WT and KO cells both before and after UVB exposure, potentially due to the underlying complexity of *AHR* regulation (**Supplementary Fig. 9**). Nonetheless, we investigated melanocyte growth phenotypes in these clones. Notably, BrdU staining and flow cytometry of wild-type and knockout clones indicated that wild-type clones had a larger proportion of BrdU-positive cells (range 42.8% to 48.6%) compared to knock-out clones (range 22.9% to 33.2%), with the heterozygous knockout clone in between (range 33.4% to 39.3%; **Supplementary Fig. 10a**). This difference of proliferation between rs117132860-WT and KO cells is confirmed by crystal violet staining of cells collected at different times after seeded at the same number (**Supplementary Fig. 10b**, WT versus KO clones, D4, *P* = 0.00031 for the representative experiment shown; *P* = 1.9 x 10^-5^ for 3 experiments combined, two-tailed paired Student’s T-test.). We also investigated cellular response to UVB exposure in these cells. At 72 hours after being treated with 13mJ/cm^2^ UVB, both WT and KO cells were cell cycle arrested reflected by smaller cell numbers and the larger and flatter cell image via crystal violet staining (**Fig. 6b, Supplementary Fig. 11**). At day 7 following UVB exposure, while many WT cells survived the UVB and went back to normal cell shape and cycle (“smaller cells” in **Fig. 6b** and **Supplementary Fig. 11**), most KO cells remained arrested (large, flat cells), indicating a clearly different cellular response to UVB treatment between WT and KO cells. BrdU staining and flow cytometry confirmed a mixed population of growing and arrested cells at day 7 following UVB exposure, with significantly more cells remaining arrested in KO lines (**Fig. 6c, Supplementary Fig. 12**).

**Fig. 6:**
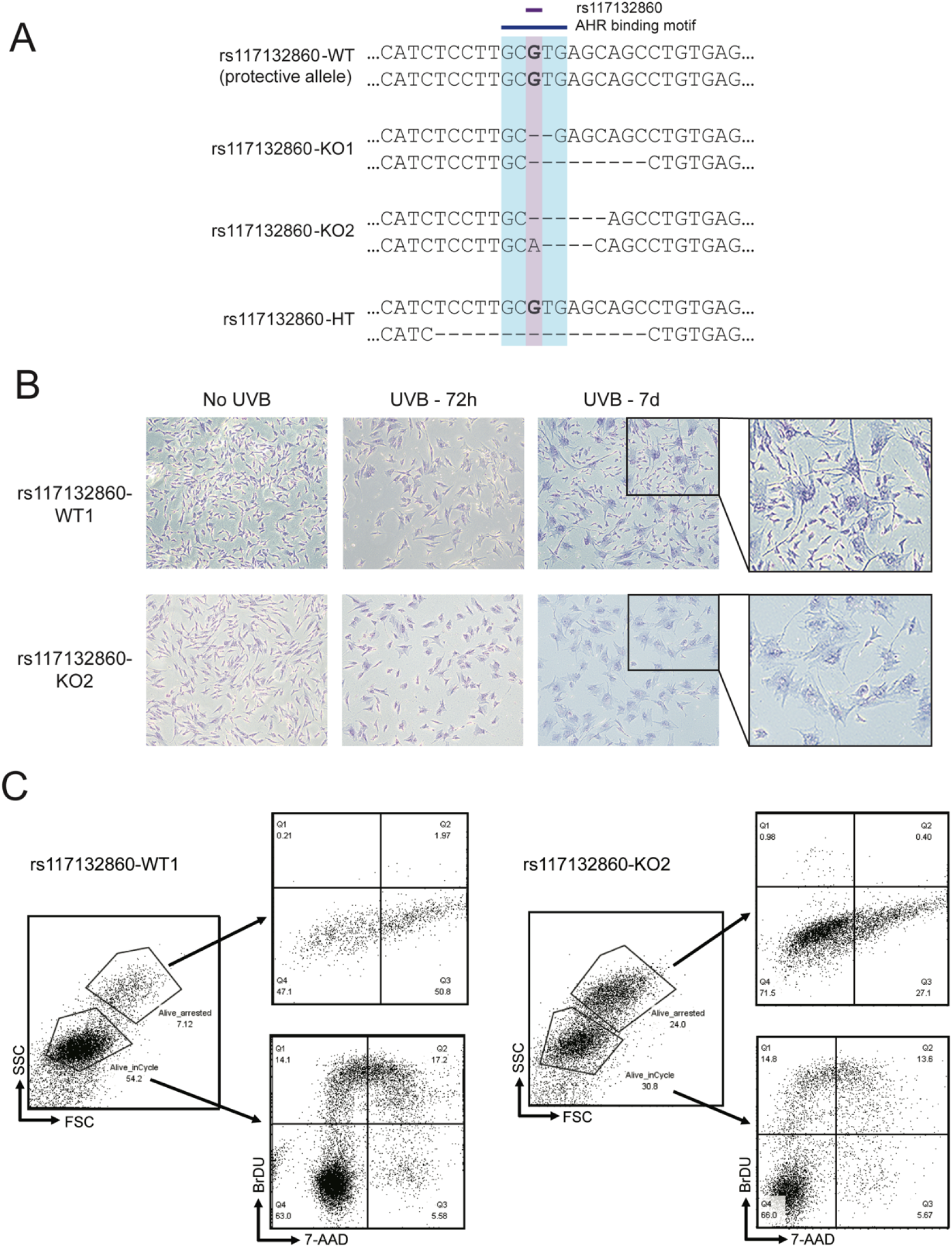
Functional characterization of CRISPR-Cas9 edited monoclonal melanocyte cell lines. (A) Genomic DNA sequences of (multiple) monoclonal cells homozygous for rs117132860-G (WT), heterozygous monoclonal cells with one wild-type allele and second copy with a 16bp deletion including the SNP (HT), and two monoclonal clones each with both alleles harboring a short deletion encompassing the AHR binding motif (KO1 and KO2). (B) Crystal Violet staining of rs117132860-WT1 and rs117132860-KO2 cells not treated with UVB (24h), 72 hours after UVB treatment, and at day 7 after UVB exposure, followed by a zoom-in image of day 7. (C) FACS analysis of the same WT and KO cells at day 7 following UVB treatment. Forward-scatter and side-scatter image indicates a mixed population with, BrdU and 7-AAD staining characterizing the cell cycle status of each population. The images shown here are from a representative experiment from three biological replicates. Additional WT and KO clones were assessed similarly.

## Discussion

Recent melanoma GWAS have successfully identified 68 independent signals at 54 genomic loci^14^. Of these, almost half are also associated with ease of tanning, reflecting the well-established interplay between pigmentation traits, UV exposure, and melanoma risk, and suggesting that environmental factors may work together with genetic variation leading to melanoma development. Here, we investigated a complex melanoma risk signal on chromosome band 7p21, fine-mapping three to four independent genetic signals, demonstrating that candidate causal variants for at least two of these physically interact with the *AHR* promoter in melanocytes. We also show that in primary melanocytes, *AHR* levels correlate with genotype at the lone candidate causal variant marking the strongest signal, rs117132860, where lower *AHR* levels are associated with the risk/poor-tanning allele. We do not identify significant correlations between rs117132860 and other nearby genes, nor do we identify chromatin loops in melanocytes to the promoters of genes other than *AHR*. Taken together, these data strongly suggest that these signals likely function via allele-specific *cis*-regulation of *AHR*.

We performed fine-mapping via conditional and joint analyses^48^ as well as using a Bayesian approach ^33,34^, as it has been suggested that Bayesian fine-mapping methods may perform better than stepwise conditional regression methods^49^. Conditional and joint analysis identified three sets of candidate causal variants between *AGR3* and *AHR* (**Fig. 1, Supplementary Figs. 1-2, Supplementary Tables 1-3**), which correlated to previously reported signals^4,14^. Conditioning on all three signals revealed a fourth marginal signal, marked by rs11766872 (*P*_conditional_ = 4.81 x 10^-6^; **Supplementary Fig. 2d, Supplementary Table 4**) approximately 400 kb from the *AHR* promoter, nearest the *SNX13* gene. Bayesian analysis using DAP-G ^33,34^, on the other hand, identified three clusters of candidate causal variants largely overlapping the first three identified by conditional analysis, with a fourth also located between *AGR3* and *AHR* (**Fig. 1, Supplementary Fig. 1, Supplementary Tables 5-6**); note the window size used for DAP-G did not include SNPs from the fourth conditional signal. Further functional studies, as well as perhaps fine-mapping using data from larger meta-analyses, will be required to better resolve the spectrum of causal variants in this region.

Still, integration of melanocyte epigenomic, melanoma functional fine-mapping (MPRA), and melanocyte Capture-C data nominates at least one strong candidate for a causal variant marking the second most significant melanoma risk signal (marked by rs73069846). Notably, of four candidate causals for this signal that directly loop to the promoter of *AHR* (rs847404, rs1721040, rs12535242, rs35785866), only rs847404 both lies within open chromatin regions in melanocytes and exhibits significant allele-specific reporter activity (rs847404, *P* = 2.62 x 10^-5^ for UACC903/HEK combined, risk allele = higher expression; **Supplementary Table 9**). MPRA data does nominate additional candidate causals for which we do not observe a direct loop between fragments containing the variant and *AHR* (rs73069846, *P* = 9.24 x 10^-13^ for UACC903/HEK cells combined; *P* = 0.0077 for UACC903 cells alone; risk allele associated with higher reporter activity; rs847377, *P* = 0.0095 for UACC903/HEK, risk allele = lower expression; **Supplementary Table 9**). Given that many restriction fragments in the broader Signal 2 region surrounding these variants were observed to associate with the *AHR* gene in melanocytes via Capture-C, these data suggest that looping between these additional candidate causals and *AHR* is plausible. Finally of note, not all candidate causal variants were assessed via MPRA or baited for Capture-C analysis, and thus we cannot rule out other variants as underlying risk in this region. Further work will be required to comprehensively assess plausible functional variants in this region.

As rs117132860 is the sole candidate causal variant associated with the lead signal at this locus, we adopted multiple approaches to characterize its potential functions in melanoma development. We used human primary melanocyte cultures as our model system to reflect tissue specific effects. Region-focused Capture-C showed a clear association between the region encompassing rs117132860 and the promoter and gene body of *AHR*, which we further corroborated by 3C mapping in multiple melanocyte cultures. eQTL analysis of 106 primary melanocyte cultures similarly nominated *AHR* as the candidate causal gene for which *cis*-regulation through rs117132860 is likely. Both luciferase reporter assay and ChIP assays demonstrated that AHR binds to its conserved binding motif containing the rs117132860-G allele and regulates expression in allele-preferential manner. As the signal marked by rs117132860 is also associated with tanning ability, cells were exposed to UVB, and we observed both increased allele-preferential binding of AHR to rs117132860-G, as well as increased ratio of expression driven from the rs117132860-G-risk versus rs117132860-A-protective alleles. Thus, our work highlights interaction between a common genetic risk variant and a well-established environmental exposure.

AHR is a well characterized pleiotropic sensor of environmental factors and has been reported to affect skin pigmentation and tanning ability after UVB exposure in a mouse knockout model, where reduced pigmentation and tanning response in knockout mice functioned through melanocyte-rather than keratinocyte-specific expression of *Ahr*^26^. Here, we were not able to test if rs117132860 is directly involved in melanocyte pigmentation due to the limitation of our cell model, as the immortalized melanocytes we used had dramatically reduced pigmentation following immortalization by hTERT with diminished expression of *TYPR1* and *TYPR2* (data not shown) relative to parental cells. Still, CRISPR-Cas9 editing of melanocytes to interrupt the AHR-binding motif harboring rs117132860-G demonstrated a clear phenotypic impact of loss of this motif in human melanocytes, including reduced proliferation as assayed by both BrdU staining as assessed by flow cytometry (**Supplementary Fig. 10a**) and crystal violet staining in a growth assay (**Supplementary Fig. 10b**), consistent with the observed negative selection of *AHR* knockout itself on cell proliferation in our CRISPR screen (**Supplementary Table 16**). This observation is consistent with a microarray analysis of cultured primary melanocytes from *Ahr*^WT^ and *Ahr^-/-^* mice, finding 3.6 to 6-fold lower expression of Endothekin-1 (*end1*) and kit ligand (*kitl*) in *Ahr^-/-^* mice, two factors involved in melanocyte proliferation ^26^. We also observe a prolonged growth arrest following UVB irradiation in rs117132860-KO cells, with a heterozygous knockout clone showing an intermediate phenotype. This raises the question of whether these arrested/senescent cells can promote secretion of an array of cytokines, chemokines and growth factors known as the senescence-associated secretory phenotype (SASP), which can create a permissive microenvironment to initiate carcinogenesis ^50,51^. While we have not investigated the possible link between rs117132860 and AHR activity induced inflammation upon UVB due to the limitations of our system, multiple studies have been reported on the role of AHR in the regulation of the immune response in the context of inflammation and cancer ^25,52,53^, as well as the potential opportunities of developing AHR targeted therapeutics. In particular, a recent report suggests a function of AHR activation on inflammation-induced melanoma cell dedifferentiation and metastases in a mouse model^44^. Taken together, these data support the notion that rs117132860 may modulate tumor risk at least in part by regulating AHR function in melanocyte proliferation and or growth arrest before and after UVB exposure, implying an additional mechanism associated with melanoma risk other than the reported effect through skin pigmentation.

Still, the response of gene targets regulated by AHR is complex, and further work will be required to better characterize the dynamics of transcriptional response to UVB irradiation. Firstly, our fine-mapping clearly shows multiple melanoma risk signals beyond rs117132860, which itself has a relatively low minor allele frequency. These data suggest multiple enhancers under genetic regulation combine to regulate AHR. Beyond this, *cis*-regulation via the enhancer harboring rs117132860 is likely to be more complex. Specifically, while the AHR/ARNT complex is generally a positive regulator of gene expression, AHR itself induced a transcriptional repressor (AHRR) which like AHR forms a complex with ARNT, binds to the consensus AHR motif, but negatively regulates expression. Further, this motif shares similarity with that bound by HIF/ARNT. In addition, given the dependency of both AHR and AHRR activity on availability of ARNT, ARNT levels and competition by other ARNT-binding transcription factors (HIF) may influence this response. Finally, our motif analysis also suggested that rs117132860-A may create a MYC binding site, and ChIP sequencing for MYC by ENCODE identified this region as bound by MYC in some cell types. Indeed, consistent with this complexity we observe several results suggesting that a simple model whereby AHR binding to rs117132860-G only upregulates *AHR* cannot fully explain short-term and long-term transcriptional dynamics upon UV exposure. While genotyping of a rs117132860 proxy SNP within the *AHR* mRNA sequence (rs17779352) after UVB exposure demonstrated higher *AHR* expression driven from the protective rs117132860-G/rs17779352-C allele in cells heterozygous for both SNPs, we failed to observe such a difference following TCDD treatment at multiple timepoints. Further, we did not observe a consistent difference in *AHR* transcriptional dynamics following UVB in rs117132860-KO (or -HT) cells. Further work will be required to characterize the dynamics of various rs117132860 allele-specific binding proteins both under normal growth conditions, as well as in response to UVB.

Finally, our findings linking the 7p21 melanoma susceptibility region to *AHR* suggests that signaling through the AHR pathway could be more broadly important in melanoma risk. Notably, multiple additional melanoma risk loci harbor genes within this pathway. Specifically, the risk locus on chromosome band lq21.3 (marked by rs8444) harbors the gene encoding ARNT, the binding partner of AHR itself on which the activity of AHR and AHRR depend. Notably, a transcriptome-wide association study using the same GWAS summary statistics found imputed melanocyte-specific *ARNT* expression to be negatively correlated with melanoma risk^14^, while TWAS of tanning found a positive correlation between predicted *ARNT* levels and ability to tan ^41^. Furthermore, a missense variant in *CYP1B1* (rs1800440), a direct transcriptional target of AHR, has been reported to be associated with both melanoma risk^4,14^ and keratinocyte skin cancers^54^. Thus, our work highlights a potential role for the larger AHR pathway in melanoma risk.

## Methods

### Fine mapping

For fine-mapping, we used melanoma GWAS summary data derived from both confirmed, as well as self-reported melanoma cases from 23andMe and UK Biobank (UKBB), and controls as previously described [14]; all participants provided informed consent and participation was IRB approved. Participants from 23andMe provided informed consent and participated in the research online, under a protocol approved by the external AAHRPP-accredited IRB, Ethical & Independent Review Services (E&I Review). Conditional- and joint-analyses of summary GWAS meta-analysis data were performed as previously described [14] using genome-wide complex trait analysis (GCTA, v.1.26.0) [48] to identify independently associated variants. To ensure we were only detecting completely independent SNPs, the collinearity threshold (--cojo-collinear) was set to *R*^2^⍰=⍰0.05. Conditional analyses of summary data in GCTA were calculated using a reference population of 5,000 individuals selected randomly from the portion of the UK Biobank population determined to be European by PCA (LDEUR). Variants were converted to best-guess genotype (threshold 0.3). Best-guess data were cleaned for missingness >3%, HWE *P*⍰<⍰1⍰×⍰^-6^ and MAF⍰<⍰0.001. Individual credible sets for each of the three association signals in the *AHR* region were identified by conditioning on the remaining two lead SNPs reported by Landi and colleagues across a 4 Mb region centered on rs117132860^14^; for each, we subsequently retained as potential causal variants with a log-likelihood ratio less than 1:100 based on conditional P-values. We further fine-mapped this region using DAPG ^33,34^. Briefly, DAP-G (version 1.0) analyses used a window size of 500 Kb centered on rs117132860 and allowing for a maximum number of 4 causals. The pairwise LD between all SNPs in each window was computed using the 1000 Genomes phase 3 EUR data using PLINK version 1.9 and R version 4.0.2.

### Cell culture

Melanoma cell lines were grown in RPMI1640 medium containing 10% FBS, 20 mM HEPES, and Amphotericin B/penicillin/streptomycin. We obtained frozen aliquots of melanocytes isolated from foreskin healthy newborn males, mainly of European descent, following an established protocol^55^ from the SPORE in Skin Cancer Specimen Resource Core at Yale University. Cells were either grown in Dermal Cell Basal Medium (ATCC PCS-200-030) supplemented with Melanocyte Growth Kit (ATCC PCS-200-041) and 1% amphotericin B/penicillin/streptomycin (120-096-711, Quality Biological) for eQTL and Capture-C analysis, or alternatively in M254 (Invitrogen, M254500) supplemented with HMGS-2 (Invitrogen, S0165) for all other experiments, grown at 37°C with 5% CO_2_. All cells tested negative for mycoplasma contamination using MycoAlert PLUS mycoplasma detection kit (LT07-710, Lonza).

### Massively Parallel Reporter Assay Data

Data were generated and analyzed as previously described ^39^. Data are presented for MPRA runs using UACC903 and HEK293FT cells analyzed jointly, as well as for UACC903 cells alone.

### Luciferase reporter assays

Luciferase constructs were generated to include the DHS region encompassing rs117132860 (375 bp; chr7:17134641-17135015). Sequences encompassing each variant were synthesized via GeneArt service from Invitrogen based on the genomic sequence of HapMap CEU panel samples; sequences are listed in **Supplementary Table 17**. Sequences were then cloned into the pGL4.23 vector using BglII and XhoI sequence overhangs. Sequence-verified pGL4.23 constructs were then co-transfected with pGL4.74 (Renilla luciferase) into the melanoma cell line UACC 903 and multiple human melanocyte cultures (c23, c197 and c87) using Lipofectamine 2000 reagent (Thermo Fisher) for UACC903 or electroporation with the Lonza Amaxa P2 kit and the CA-137 protocol on the 4D-Nucleofector system for primary melanocytes. Cells were collected 24h after transfection, and luciferase activity was measured using the Dual-Luciferase Reporter System (Promega) on the GLOMAX Multi Detection system (Promega). All experiments were performed in at least three biological replicates in sets of 4 replicates.

### eQTL analysis

Melanocyte eQTL were generated and analyzed as previously described ^40^ (dbGAP phs001500.v1.p1). Genes within the TAD were considered expressed in melanocytes based on RSEM reading with subsequent Bonferroni correction based on the number of expressed genes in the TAD (n=3; 0.05/3).

### ATAC-seq library generation and peak calls

30K-50K primary melanocytes were lysed with cold lysis buffer (10⍰mM Tris-HCl, pH 7.4, 10⍰mM NaCl, 3⍰mM MgCl2, 0.1% IGEPAL CA-630), and centrifuged to get nuclei. The nuclei were resuspended in the transposition reaction mix (2x TD Buffer (Illumina Cat #FC-121-1030, Nextera), 2.5⍰μl Tn5 Transposase (Illumina Cat #FC-121-1030, Nextera) and Nuclease Free H_2_O) on ice and then incubated for 30⍰min at 37°. The transposed DNA was then purified using the MinElute Kit (Qiagen), and PCR amplified using Nextera primers for 12 cycles to generate each library. The PCR reaction was subsequently cleaned up using AMPureXP beads (Agencourt) and libraries were paired-end sequenced on an Illumina HiSeq 4000 (100 bp read length) and the Illumina NovaSeq platform. 15 ATAC-seq libraries were made from five independent primary melanocyte cultures (C56, C140, C205, C24, and C27) with three replicates for each. ATAC sequencing data analysis was done using the ENCODE ATAC-seq pipeline version 1.6.1 (https://www.encodeproiect.org/atac-seq/). ATAC sequencing reads from independent replicates of each of the five melanocyte cultures (3 replicates per culture) were merged together. The five melanocyte cultures were treated as biological replicates and their sequencing reads were aligned to hg19 genome using bowtie2 ^56,57^, and duplicate reads were removed. Two pseudo replicates were generated by random sampling of reads from pooled biological replicates. ATAC peaks were called using MACSv2 peak caller (2.1.0)^58^ (P< 0.01) and the ENCODE blacklist regions were removed during the peak calling process^59^. Peaks were called from individual biological replicates, pooled data of biological replicates, and pooled data of pseudo replicates. Irreproducible Discovery Rate (IDR) was calculated to find peaks that are reproducible and rank consistently across individual replicates and pooled replicate data. Peaks that overlapped between individual replicates and pooled replicate data with IDR < 0.05 were selected. This peak set is also known as the conservative IDR set. The ATAC-seq peaks that overlapped between the pseudo replicates and pooled replicate data with IDR<0.05 were selected and referred to as the Pseudo IDR Set (PIS). The number of peaks between conservative IDR set and PIS were compared and the maximum of the two were selected as the optimal IDR set. The conservative IDR and optimal IDR sets passed the reproducibility test as they met the conditions of Self-consistency Ratio <2 AND Rescue Ratio < 2.

Furthermore, the peaks from both sets between the AHR Topologically Associating Domain region (chr7: 16720000-17720000) were analyzed and visualized on the UCSC Genome Browser^60^ or WashU Epigenome Browser^61^.

### Capture-C

Custom capture baits were designed by Arima Genomics (San Diego, CA) using an Agilent Sure Select library design targeting specific restriction fragments encompassing two genome-wide significant 7p21 signals from single-SNP analyses ^14^. The probe library was synthesized by Agilent. Regions for bait design were determined for each locus by identifying the set of SNPs with a log-likelihood ratio <100 relative to the leading SNP of a given GWAS signal. For the *AHR* locus, we covered signals for both genome-wide significant signals defined by Landi and colleagues ^14^ (chr7:17134618-17135137 and chr7:16966279-17005842); baited fragments are listed in **Supplementary Table 10**. Capture-C libraries were made using the Arima HiC kit (Arima Genomics) and the KAPA HyperPrep kit (KAPA Biosystems) following the manufacturer’s protocol. Briefly, 2-4 million cells were crosslinked, enzyme digested, and ligated. The ligated DNA was reverse-crosslinked, fragmented by sonication, and size-selected for adaptor ligation and library amplification. The HiC library was then hybridized with the custom capture baits and captured by the SureSelect XT HS and XT low input library preparation kit for ILM (Agilent). 15 Capture C libraries were made from 5 human primary melanocyte cultures (C56, C140, C205, C24, and C27) with 3 biological replicates for each. The captured libraries were pooled and sequenced using an Illumina Novaseq, with one run on SP and second run on S1 flowcell, generating ~5.7 billion paired-end reads with 150bp read length. Paired-end sequencing reads from biological replicates were pre-processed using the HiCUP pipeline^62^ and aligned to human genome version 19 using bowtie2 ^56,57^. Chromatin interaction loops were detected at one and four fragment resolutions using CHiCAGO pipeline version 1.16.0^42^. Default parameters were used for one fragment analysis except for minFragLen, maxFragLen, binsize, maxLBrownEst which were set to 75, 1200, 2000, and 150000 respectively. Four fragment resolution was created using artificial .baitmap and .rmap files where four consecutive restriction digestion fragments were grouped into one fragment. Four fragment analysis was conducted using default parameters except for minFragLen, maxFragLen, binsize, maxLBrownEst which were set to 150, 5000, 8000, and 600000 respectively. Chromatin interactions with CHiCAGO scores ≥ 5 were considered high-confidence interactions and were further analyzed. The output file was generated using long range interaction format and was used for visualization on the WashU Epigenome Browser^61^.

### Chromatin Confirmation Capture (3C)

3C assays were done based on protocol from Dekker Lab^63^. RP11-317K18 and RP11-594C23 BAC clones were purchased from BACPAC Resources Center and purified using Qiagen large-construct Maxi kit (#12462) to cover the genomic region between rs117132860 and the *AHR* gene to be mapped (RP11-317K18, chr7:17121094-17313112; RP11-594C23, chr7:17306808-17483385). BAC libraries were made by HindIII digestion of BAC plasmids, followed by ligation and DNA purification. To generate melanocyte 3C libraries, ~20 million cells were fixed and lysed followed by HindIII digestion and ligation. Both melanocyte 3C library and BAC libraries were amplified with a Taqman assay containing a primer localized to various regions between rs117132860 and *AHR* gene and a fixed primer near rs117132860 plus a FAM^™^ labelled probe annealing to 3’ of the fixed primer (primer sequences are listed in **Supplementary Table 18**). PCR cycle conditions were as follows: 50C, 2 min; 95C, 10 min; [95C, 15s; 60C 1 min]_X40_. Amplification for different primer pairs from the melanocyte 3C library was normalized to that of BAC libraries, reflecting the potential of chromatin interaction.

### Chromatin Immunoprecipitation

Primary melanocytes were fixed with 1% formaldehyde when ~85% confluent, following the instructions of Active Motif ChIP-IT high sensitivity kit. 7.5⍰x⍰10^6^ cells were then homogenized and sheared by sonication using a Bioruptor (Diagenode) at high setting for 15⍰min, with 20s on and 30s off cycles. 5ug sheared chromatin was used for each immunoprecipitation with antibodies against AHR (Cat#83200s, Cell Signaling), or normal rabbit IgG (Cat# ab37415, Abcam) following the manufacturer’s instructions. Purified antibody pulled-down DNA or input DNA was assayed by SYBR Green qPCR for enrichment of target sites using primers listed in (**Supplementary Table 19**). Relative quantity of each sample was derived from a standard curve of each primer set and normalized to 1/1000 input DNA. For rs117132860 genotyping, input DNA or genomic DNA from each cell line, and DNA pulled down using an anti-AHR antibody from melanocytes heterozygous for rs117132860 (c87, c197 and c262) was used as template DNA for a Taqman genotyping assay (Assay ID: ANU699F, Thermo Scientific). All experiments were performed for at least three biological replicates.

### UVB and TCDD treatment, AHR qRT-PCR and western blotting

Human primary melanocytes were treated with 10nM TCDD dissolved in toluene (Sigma, Cat#48599) for 1-24hrs. Toluene treated cells served as non-treated control. Cells were exposed with UVB (312nm) for 10s from a Spectronics ENB-280C Handheld UV Lamp. The UVB dosage was measured to be 13.2mJ/cm^2^ by a radiometer. Cells were lysed with Trizol (Thermo Scientific) for RNA prep or with RIPA buffer (Thermo Scientific) for cell extract prep. For assessment of *AHR* mRNA levels, we purified total RNA using the Qiagen RNeasy Mini kit (Cat#74106) and made cDNA using iscript cDNA synthesis kit from Biorad (Cat#1725036), followed by a Taqman assay to detect *AHR* transcription (Hs00169233-m1 *for AHR* expression and Hs0442063-g1 for *GAPDH;* Thermo Scientific). For protein analysis, whole cell extracts were subjected to water bath sonication and samples were resolved by 4–12% Bis-Tris ready gel (Invitrogen) electrophoresis. The primary antibodies used were rabbit antibody to AHR (Cat#83200s, Cell Signaling) and mouse antibody to β-actin (A5316, Sigma).

### AHR allele-specific expression

Total RNA was isolated using a RNAeasy Mini kit (217004, Qiagen). cDNA was synthesized from total RNA using iScript Advanced cDNA Synthesis Kit (Bio-Rad). Genomic DNA and cDNA were then genotyped for rs17779352 using a custom Taqman genotyping assay for rs17779352 (ANCE9Z2, Thermo Scientific; primers and probes listed in **Supplementary Table 19**) recognizing both genomic DNA and cDNA (ENST00000543692). The AHR expression ratio from the risk rs117132860-G/rs17779352-C allele relative to the risk rs117132860-A/rs17779352-T allele were calculated from dRn value based on the linkage between rs117132860 and rs17779352. All experiments were performed in at least three biological replicates in sets of triplicates.

### Gene-based CRISPR-Cas9 knockout screen

Target genes were predominantly selected from genome-wide significant melanoma risk loci^14^. Up to 10 guide RNAs per gene were designed against the list of target genes. First, sequences were drawn from pre-existing genome wide libraries ^64,65^. The remaining sequences were obtained from a published computationally predicted list^66^. 200 negative control sequences were obtained from the GeCKO V2 library^67^. Oligonucleotides, with flanking sequences for PCR sub-pool amplification and isothermal assembly, were synthesized by Twist Biosciences (**Supplementary Table 14**). Primers for sub-pool amplification (**Supplementary Table 19**) were obtained from a previously published list of sequences orthogonal to the human genome^68^. Sub-pools were PCR amplified using quantitative PCR to minimize amplification bias and PCR products were then gel extracted from a 2% E-gel EX. 5 ng of gel extracted PCR product was then assembled with 50 ng of BsmBI digested lentiGuide-Puro (PMID: 25075903) using the NEB HiFi Builder. lentiGuide-Puro was a gift from Feng Zhang (Addgene plasmid # 52963; http://n2t.net/addgene:52963; RRID:Addgene_52963).

Assemblies were then purified using the Zymo Clean and Concentrate 5 and eluted in 10 uL of elution buffer. 5 uL of eluted product was then electroporated into Lucigen Endura electrocompetent cells using the BioRad Gene Pulser. Cells were recovered with SOC and plated across 5 - 15 cm plate and grown at 30C. ~18hrs later, colonies were immersed in LB, scraped and pelleted, and plasmid DNA was extracted using the Qiagen Plasmid Plus Maxiprep kit. For production of pooled lentivirus expressing gRNA, the pooled plasmids were co-transfected into HEK293 cells with the pSPAX2, pMD2-G, and pCAG4-RTR2 packaging vectors. Virus was collected 2 days after transfection and concentrated by Vivaspin. Virus titre was measured by HIV-1 p24 Antigen Elisa Kit by ZaptoMetrix Corporation. The TERT-immortalized human melanocyte cell C283T was infected with pCW-Cas9-Blast from Addgene followed by introduction of lentiGuide-Puro (Addgene) expressing pooled gRNAs. Infection rate was controlled at ~0.3 to allow no more than one copy of sgRNA in each cell. After drug selection and cell recovery (~ day 7 after infection; NoDOX;), a fraction of cells was collected, and the remaining cells were treated with 0.5ug/ml dox to induce Cas9 expression. Cells were then collected at roughly day 14 (DOX_T1), day 21 (DOX_T2) and day 28 (DOX_T3) after infection (**Supplementary Table 15**) and DNA was purified for sequencing by MiSeq. The raw sequences were aligned conservatively (only perfect matches counted) via MAGeCK and the normalized read counts for each sgRNA from samples before and after dox treatment (three biological replicates; Exp1, Exp2, Exp3) were compared and analyzed by MAGeCK together in order to rank both positively and negatively selected genes. Genes and gRNAs are supplied in **Supplementary Table 14**.

### CRISPR-Cas9 editing of the rs117132860 AHR-binding motif

We designed a sgRNA targeted to rs117132860 (**Supplementary Table 19**) to introduce small deletions into the AHR-binding motif encompassing rs117132860. The sgRNA was cloned and expressed in LentiGuide-Puro (Addgene, Cat#52963). A Cas9-expressing lentiviral vector pCW-Cas9-Blast was virally introduced into an TERT-immortalized human melanocyte culture, C283-T, followed by further introduction of either the plasmid containing sgRNA targeted to rs117132860 or a non-targeting sgRNA (Addgene, Cat#80226). Following selection, we isolated monoclonal cell lines from the mixed population through limited dilution. For candidate clones grown from single cells, we sequenced the genomic region around rs117132860 and deconvoluted the sequences to identify potential cell clones with deletion/mutation regarding to AHR-binding motif around rs117132860 (Mutation Surveyor, Soft Genetics). We identified 2 clones with deletion/mutation of both alleles of the AHR-binding motif around rs117132860 (KO1 and KO2), one clone with one copy of wild type sequence and one copy with deletion of AHR binding motif (HT), and 2 clones with no change of the sequence around the SNP (WT).

### Cell cycle and proliferation assays

Cell proliferation was assayed using a BrdU flow kit (BD Pharmingen) according to the manufacturer’s protocol. Briefly, human melanocytes were labeled with 10 μM BrdU for 3 hours before they were fixed, permeabilized, and subjected to DNase I treatment. Cells were then stained with FITC-conjugated antibody to BrdU and 7-AAD, followed by flow cytometry analysis using a BD Accuri instrument (BD Pharmingen). For crystal violet staining, cells were seeded at equal numbers and subjected to UVB exposure before being fixed and stained with crystal violet.

### Statistical analyses

All cell-based experiments were repeated at least three times with multiple cell cultures. When a representative set is shown, replicate experiments displayed similar but not identical patterns. For all plots, mean and SEM are shown, except for violin plots where individual data points are shown with the median or mean, range (maximum and minimum), and 25th and 75th percentiles (where applicable). The statistical method, number of data points, and number and type of replicates are indicated in each Fig. legend.

### Predicted transcription factor binding analysis

To predict the effects of candidate causal variants on potential transcription factor binding sites, we used motifbreakR^38^ version 2.4.0 on a total of 56 fine-mapped CCVs. Analysis used dbSNP144 for GRCh37, the UCSC genome build hg19 for the reference sequence, and 2,817 position frequency matrices from ENCODE-motif, FactorBook, HOCOMOCO, and HOMER (provided through the ‘MotifDb’ package, https://bioconductor.org/packages/release/bioc/html/MotifDb.html), and using the default significance threshold of 1.0 x 10^-4^. Six variants were dropped from analysis (rs35785866, rs200020478, rs199662382, rs10532327, rs367629062, and rs10589929) as they were not included in dbSNP144/GRCh37. All motifbreakR analyses were run using R version 4.0.3.

## Data Availability

Conditional fine-mapping data is available in Supplementary Tables 1-4; Bayesian fine-mapping results in Supplementary Tables 6-7. Melanocyte ATAC seq peaks for the region are available in Supplementary Table 7. MotifbreakR analysis is provided in Supplementary Table 8. Data from the 2020 melanoma GWAS meta-analysis performed by Landi and colleagues was obtained from dbGaP (phs001868.v1.p1), with the exclusion of self-reported data from 23andMe and UK Biobank. The full GWAS summary statistics for the 23andMe discovery data set will be made available through 23andMe to qualified researchers under an agreement with 23andMe that protects the privacy of the 23andMe participants. Please visit https://research.23andme.com/collaborate/#dataset-access/ for more information and to apply to access the data. Summary data from the remaining self-reported cases are available from the corresponding authors of that manuscript (Matthew Law; Mark Iles; and Maria Teresa Landi). MPRA data are available in the NCBI Gene Expression Omnibus (https://www.ncbi.nlm.nih.gov/geo/) as a SuperSeries under the accession number GSE129250. Restriction fragments baited for region-specific capture-C assays are provided in Supplementary Table 10, with loops called by ChICAGO in Supplementary Tables 11-12. gRNAs for the pooled gene-based CRISPR screen are provided in Supplementary Table 14, with screen read counts for each and MaGeCK output available in Supplementary Tables 15 and 16, respectively. Luciferase assay fragments and other primers and gRNAs are listed in Supplementary Tables 17-19.

## Acknowledgements

We would like to thank members at the National Cancer Institute Cancer Genomics Research Laboratory (CGR) for help with sequencing efforts, and L. Amundadottir from National Cancer Institute, Laboratory of Translational Genomics for helpful discussions This work has been supported by the Intramural Research Program (IRP) of the National Cancer Institute, US National Institutes of Health, including through an NCI FLEX award to G.M. and K.M.B. M.M.I. is supported by Cancer Research UK (Programme award C588/A19167) and by the National Institutes of Health (R01 CA83115), with GWAS genotyping services provided by the Center for Inherited Disease Research (CIDR); CIDR is fully funded through a federal contract from the National Institutes of Health to The Johns Hopkins University, contract number HHSN268201100011I. M.H.L acknowledges funding support from the Australian National Health and Medical Research Council (NHMRC; APP1129822, APP1123248) and Worldwide Cancer Research (WCR16-101). We also thank all the cohorts, funders, and investigators who contributed to the melanoma GWAS, as originally acknowledged by Landi, 23andMe, and colleagues ^14^, from which data was used towards fine-mapping. S.F.A. Grant is supported by NIH R01 HG010067 and the Daniel B. Burke Endowed Chair for Diabetes Research. The content of this publication does not necessarily reflect the views or policies of the US Department of Health and Human Services, nor does mention of trade names, commercial products, or organizations imply endorsement by the US government.

## Author contributions

K.M.B and M.X. conceived and planned the study. M.H.L. performed conditional fine-mapping and H.S., R.T., and M.X. performed Bayesian fine-mapping, both with input from M.T.L, M.M.I, M.H.L., and A.M.G. Melanocyte ATAC-seq data was generated by M.E.G., A.D.W., A.C., and S.F.A.G., and analyzed by R.T. M.X., T.M., L.J., K.J., S.J.C. and H.H. designed and performed Capture-C experiments; M.X., R.T., and H.S. performed Capture-C data analysis. J.C., T.Z., and M.A.K. performed QTL analyses. J.C. and T.Z. performed MPRA analyses. M.X. and L.M. performed luciferase assays, ChIP, and exposure analyses of primary melanocytes. K.M.B., G.M., M.X., R.C., H.T.M., and P.B. designed and performed gene-based CRISPR screens in melanocytes. M.X., R.C., and T.Z. designed and performed region-based CRISPR knockouts and subsequent characterization experiments. M.X. and K.M.B. wrote the manuscript. K.M.B. and J.C. helped supervise the project.

## Ethics declarations

The authors declare no competing interests.

## Notes

### Competing Interest Statement

The authors have declared no competing interest.

