## Supplementary Figures for "A UVB-responsive common variant at chr7p21.1 confers tanning response and melanoma risk via regulation of the aryl hydrocarbon receptor gene (*AHR*)"

**Supplementary Figure 1: Fine mapping of multiple melanoma GWAS signals on chromosome band 7p21.1.** (Top) a wider view (chr7:16600000-17500000) of the same 7p21.1 locus depicted in Figure 1, including UCSC genes, imputed ChromHMM and DNaseI hypersensitivity (DHS) data from two melanocyte cultures generated by the RoadMap Project, ATAC-seq data generated from five human primary melanocyte cultures, and candidate causal SNP sets nominated by either conditional analysis or Bayesian fine-mapping (95% credible sets for each of four clusters) using DAP-G. Candidate causal variants mapping to signal/cluster 1 are in red, signal/cluster 2 in green, signal/cluster 3 in blue, and signal/cluster 4 in purple. (Bottom) a zoomed-in view of the region immediately surrounding rs117132860, showing rs117132860 mapping to open chromatin and annotated melanocyte enhancer. Genomic positions are based on hg19. For the zoomed-in region containing rs117132860 (bottom), layered 7-cell H3K27Ac, DHS clusters, and transcription factor CHIP tracks were generated by ENCODE and provided through the UCSC Genome Browser.

### Supplementary Figure 1

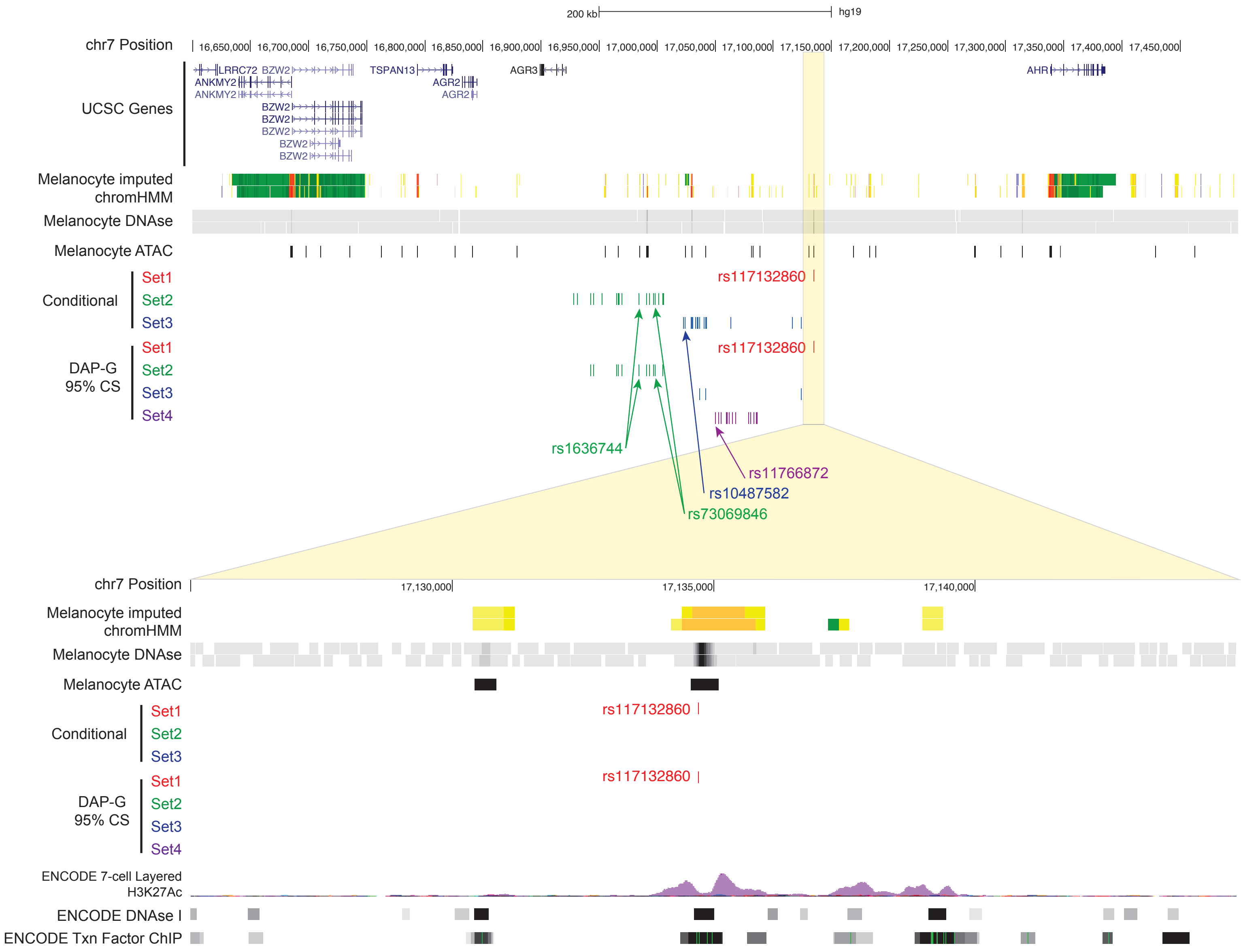

**Supplementary Figure 2: Manhattan plots of conditional association signals from the melanoma GWAS locus on chromosome band 7p21.1.** $-\log_{10}$ *P* values for SNPs at 7p21.1 when conditioned on (A) rs73069846 and rs10487582, (B) rs117132860 and rs10487582, (C) rs117132860 and rs73069846, and (D) rs117132860, rs73069846, and rs10487582. Analysis was performed using genome-wide complex trait analysis (GCTA, v.1.26.0).

Supplementary Figure 2

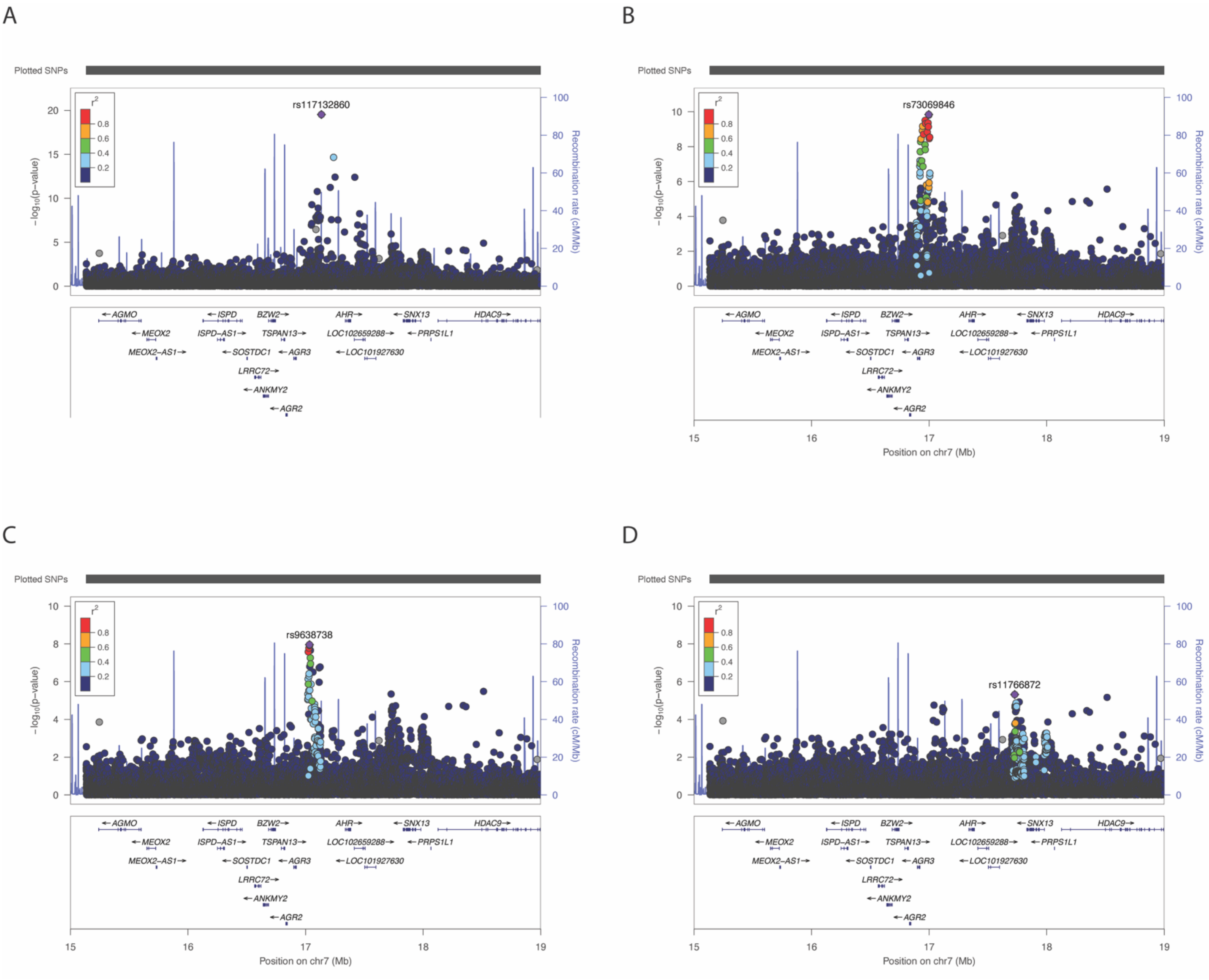

**Supplementary Figure 3: Region-specific capture-C shows chromatin looping between multiple independent 7p21 GWAS signals and the *AHR* promoter and gene body.** Significant chromatin interactions captured by capture-C between 7p21.1 melanoma risk signals and the *AHR* gene. Region capture baits are labeled in black and significant interactions are shown as purple arcs. UCSC genes, imputed ChromHMM and DNaseI hypersensitivity (DHS) data from two melanocyte cultures generated by the RoadMap Project, ATAC-seq data generated from five human primary melanocyte cultures, and candidate causal SNP sets nominated by either conditional analysis or Bayesian fine-mapping (95% credible sets for each of four clusters) using DAP-G are also shown. Loops were called from data from five distinct melanocyte cultures (3 biological replicates per culture) analyzed together in order to detect the most reproducible interactions. Baited 7p21 Signal 1 and Signal 2 regions are highlighted (yellow), both of which show interactions to the *AHR* gene (blue).

### Supplementary Figure 3

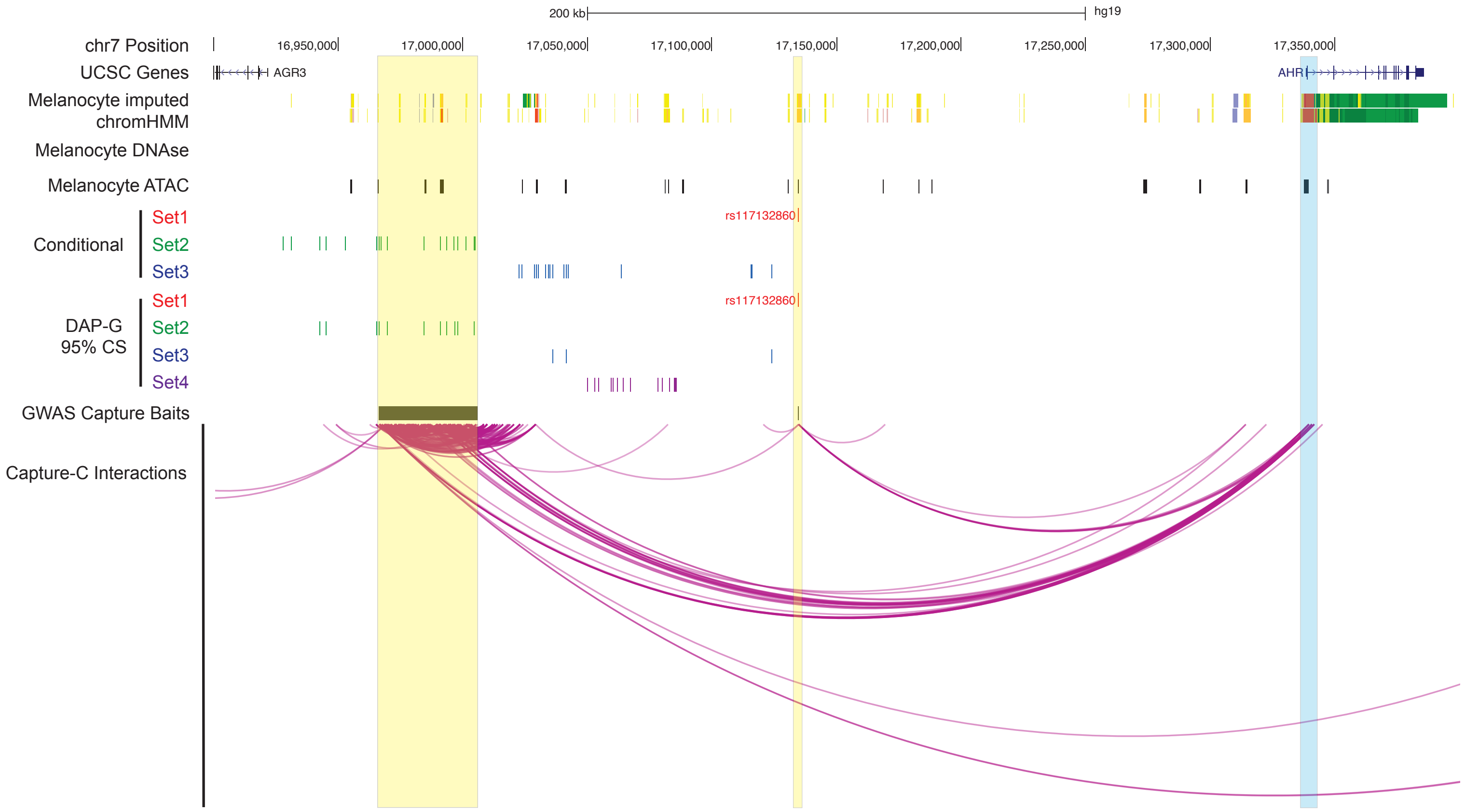

**Supplementary Figure 4: Dynamic change of *AHR* expression in additional melanocyte cultures upon TCDD and UVB exposure.** (A) *AHR* transcription normalized to *GADPH* was measured by Taqman assay in human C197 human melanocytes before and after TCDD treatment. (B) *GAPDH* normalized *AHR* transcription increased after UVB exposure in C23 and C262 human melanocytes. All experiments were done with a total of four biological replicates each, with the figure depicting one representative replicate. *P* values are calculated from a two-way paired Student's T-test and the mean with SEM is plotted (*: $p<0.05$; **: $p<0.01$;).

Supplementary Figure 4

A

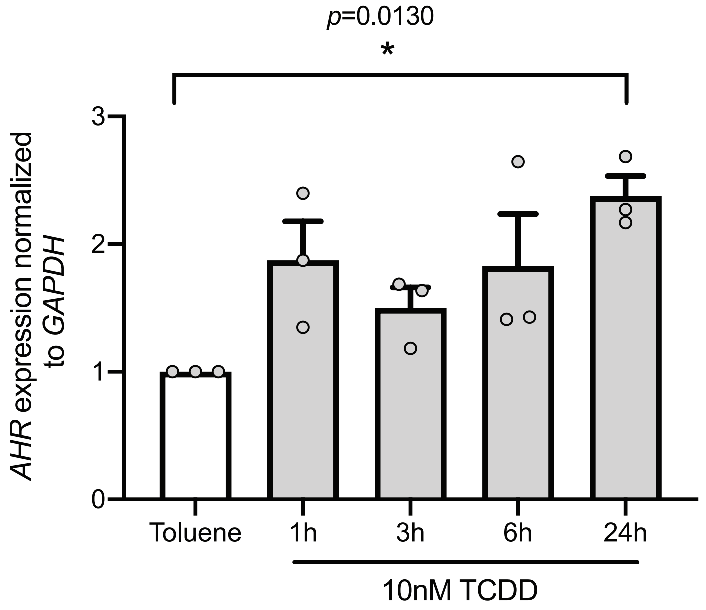


B

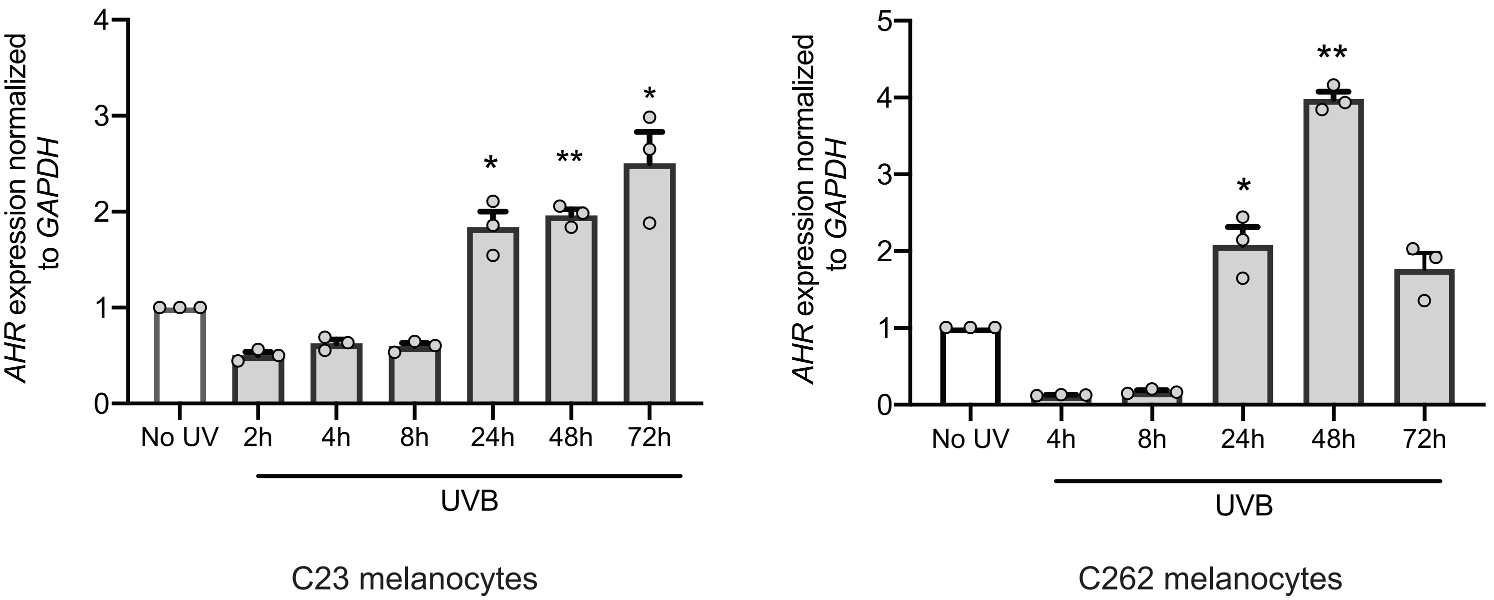

**Supplementary Figure 5: Dynamic change AHR binding to rs117132860-G in additional melanocytes culture upon TCDD and UVB exposure.** AHR binding to rs117132860 measured by CHIP assay both increased after (A) TCDD treatment and (C) UVB exposure in C197 human melanocytes. A genotyping assay using AHR ChIP DNA shows enhanced binding of AHR to the melanoma protective rs117132860-G allele both after (B) TCDD treatment and (D) UVB exposure in C197 human melanocytes. All experiments were done with a total of four biological replicates each, with the figure depicting one representative replicate. *P* values are calculated from a two-way paired Student's T-test and the mean with SEM is plotted.

Supplementary Figure 5

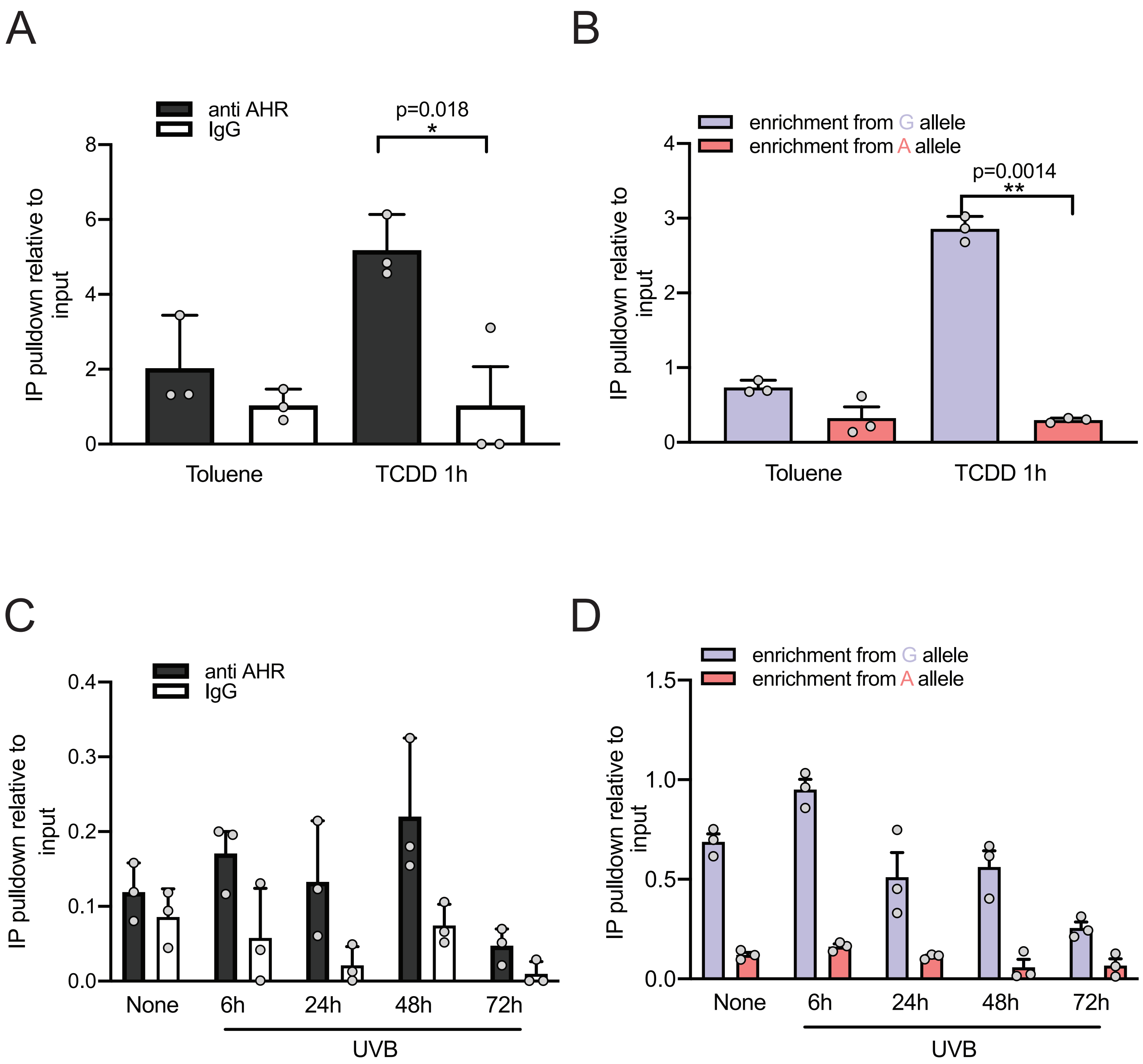

**Supplementary Figure 6: Allele-specific *AHR* expression following TCDD.** A genotyping assay of rs17779352, a proxy SNP for rs117132860 located in in the *AHR* coding sequence, indicates that the ratio of *AHR* expression from the melanoma-protective rs117132860-G/rs17779352-C allele remains unchanged relative to the rs117132860-A/rs7779352-T allele following TCDD exposure at from 1 to 24 hours in C87 human melanocytes. The proxy SNP genotyping experiment was only performed in C87 cells which is heterozygous for both rs117132860 and rs17779352; the violin plot shows the combination of 4 experiments of 3 replicates each with all points shown.

Supplementary Figure 6

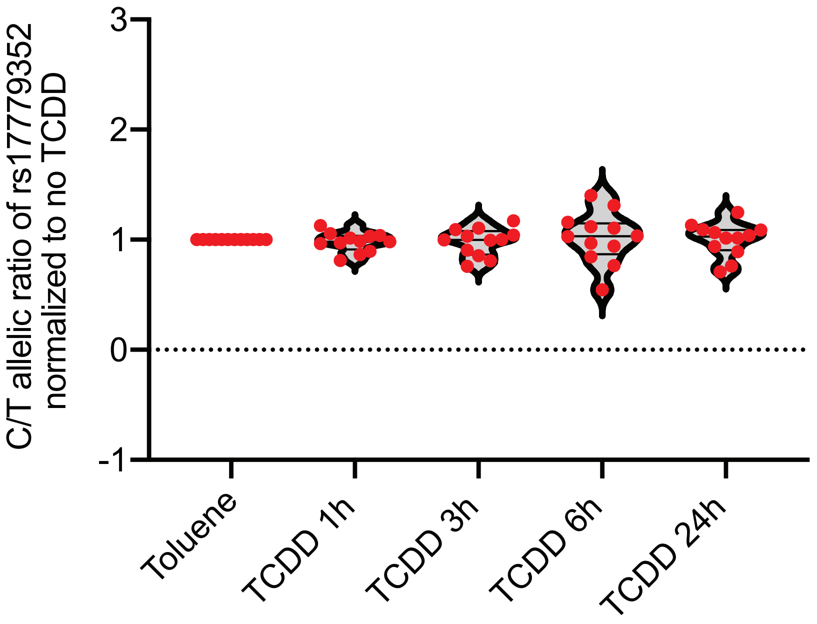

**Supplementary Figure 7: Sanger sequencing traces and deconvolution for monoclonal CRISPR-Cas9 edited rs117132860 knock out clones via Mutation Surveyor.** For each clone, the top track is the reference sequence trace, the second to top is the actual sequence trace by sanger sequencing, the third and fourth traces are the deconvoluted sequence for each copy of the gDNA, and the fifth is the shifted sequence trace of the copy containing deletion around rs117132860.

### Supplementary Figure 7

## rs117132860-KO1

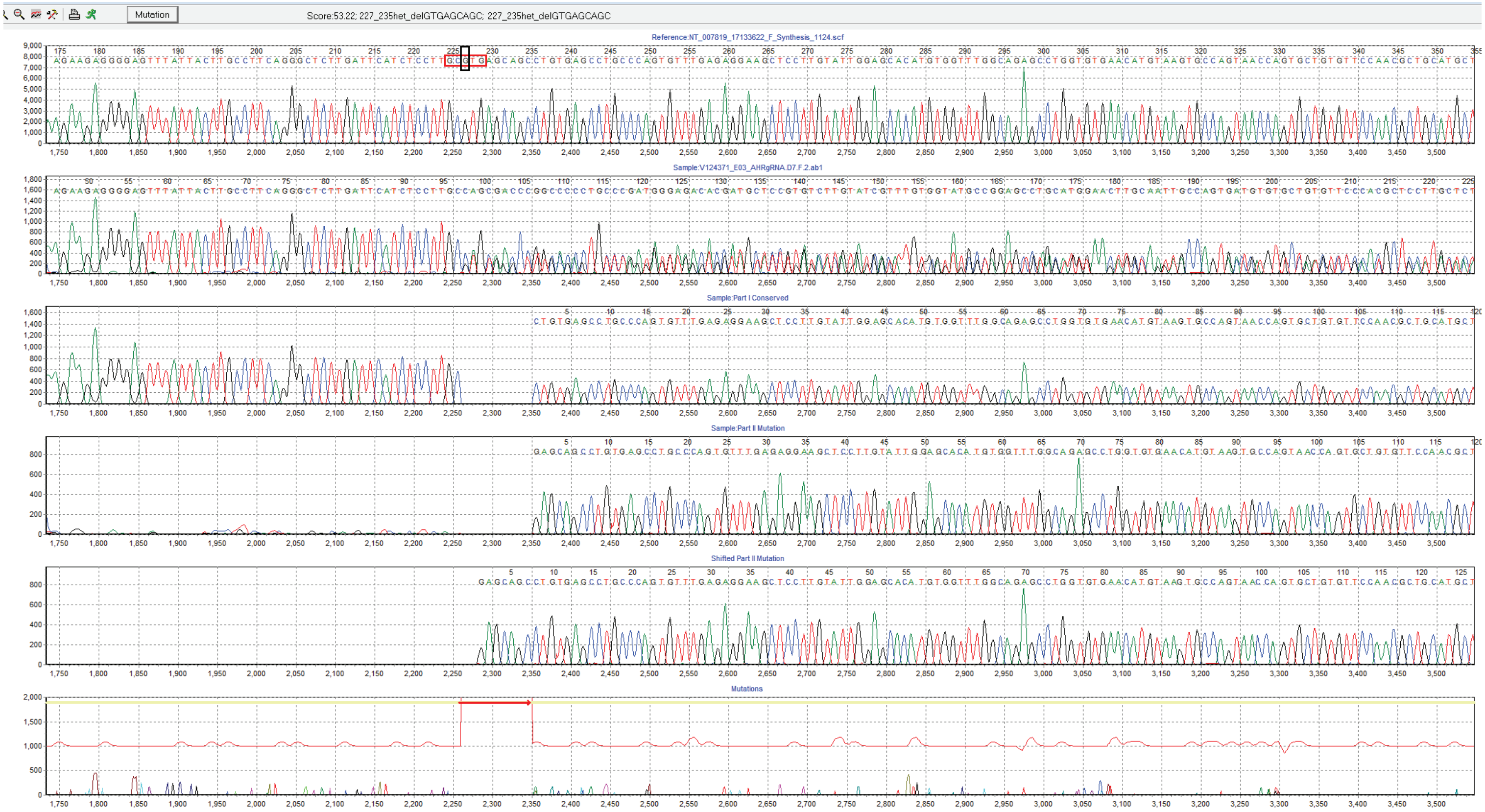


## rs117132860-KO2

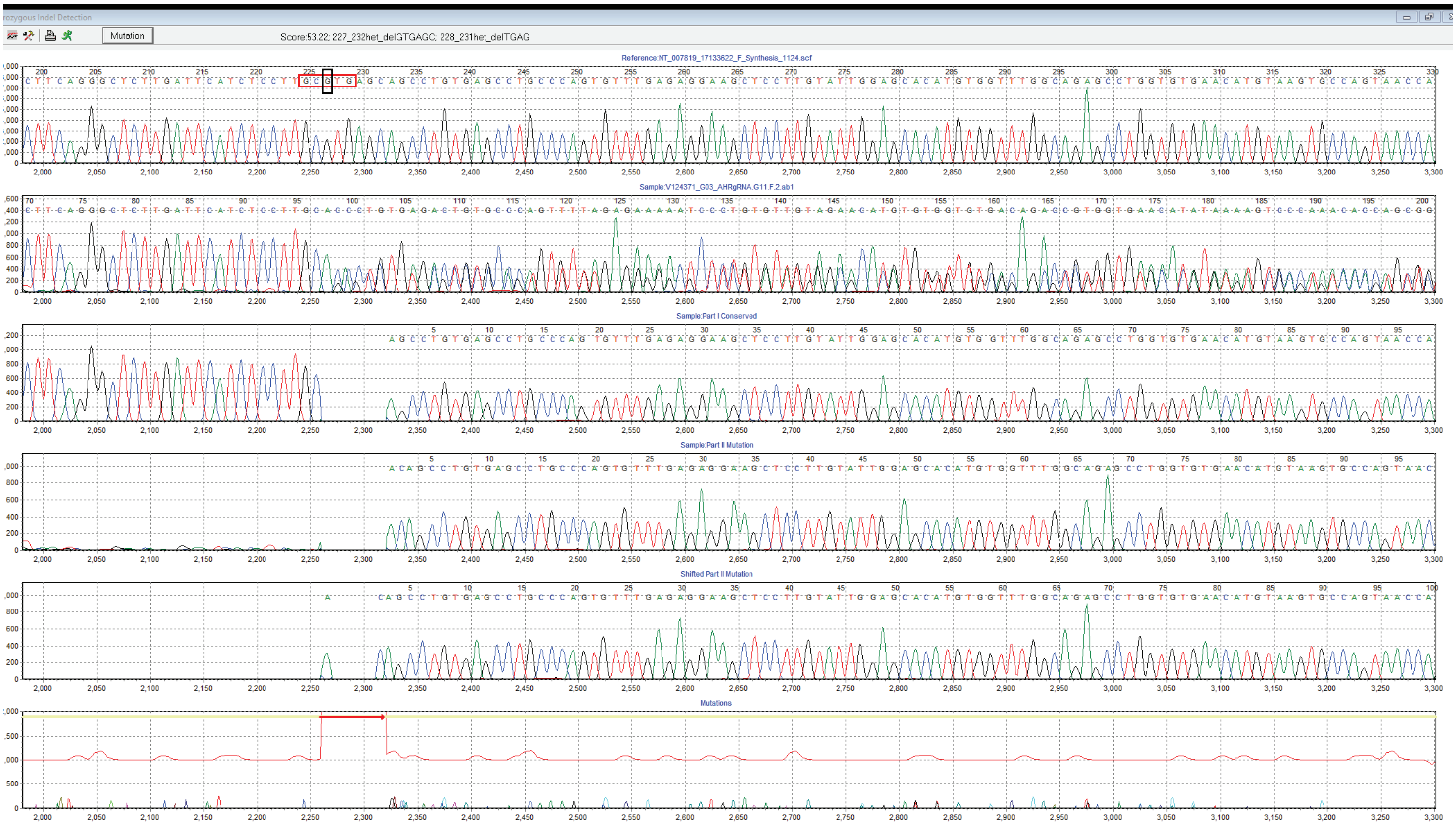

**Supplementary Figure 8: AHR binding to rs117132860 region via CHIP assay in rs117132860 knock out clones.** AHR binding to rs117132860 measured by CHIP assay increased after UVB exposure in two rs117132860-WT clones, while both rs117132860-KO and rs117132860-HT cells do not show significant AHR binding both before and after UVB exposure. A single representative experiment of two separate experiments (3 technical replicates per experiment) is shown for WT and KO clones (HT was only done once). Mean and SEM are graphed.

### Supplementary Figure 8

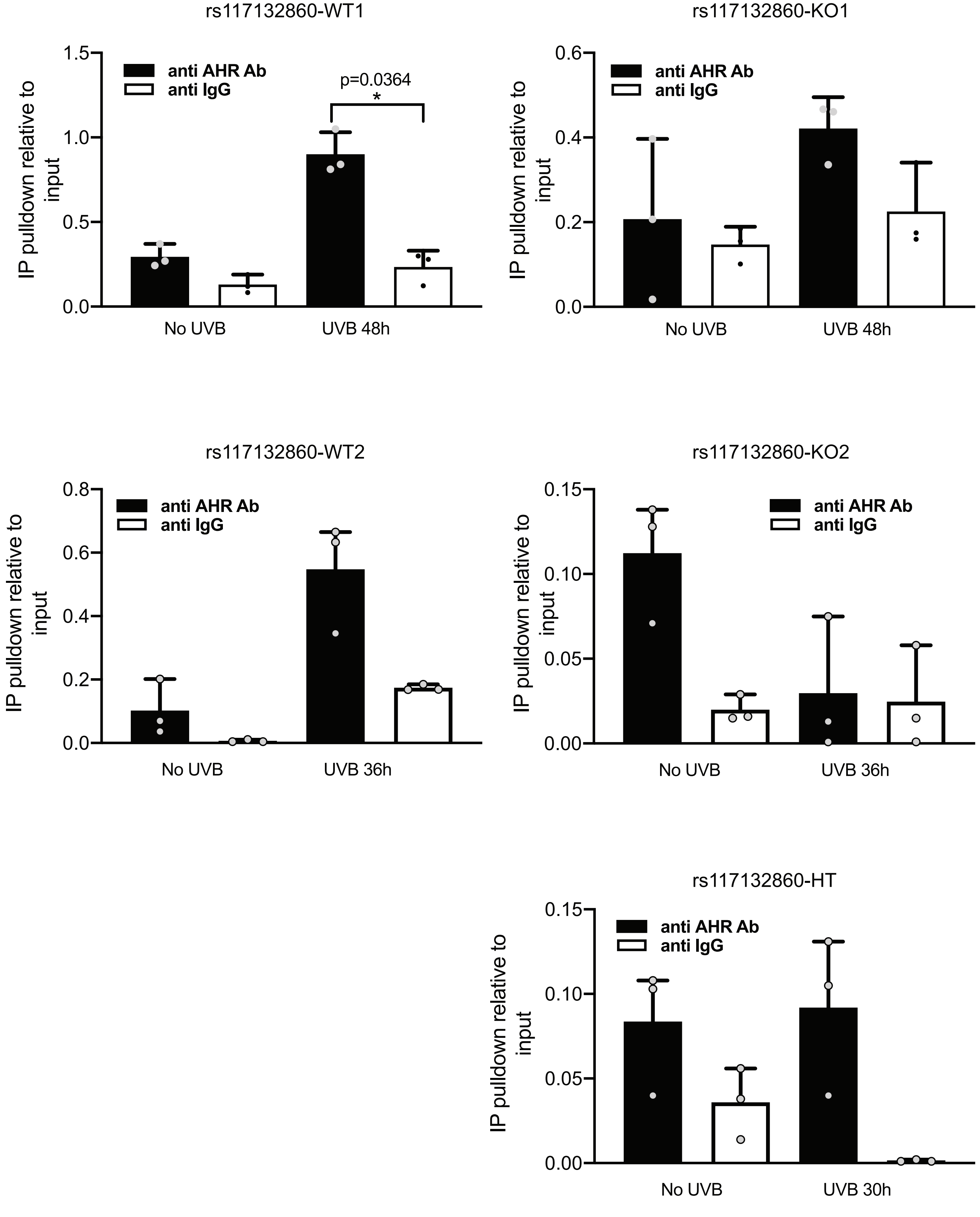

**Supplementary Figure 9: *AHR* expression in rs117132860 knock out clones with and without UVB treatment.** *AHR* expression as measured by quantitative RT-PCR for rs117132860- WT, rs117132860-KO, and rs117132860-HT clones both without and following UVB treatment. Results from four separate biological replicates are shown; AHR expression is normalized to *GAPDH*. Mean and SEM are graphed.

Supplementary Figure 9

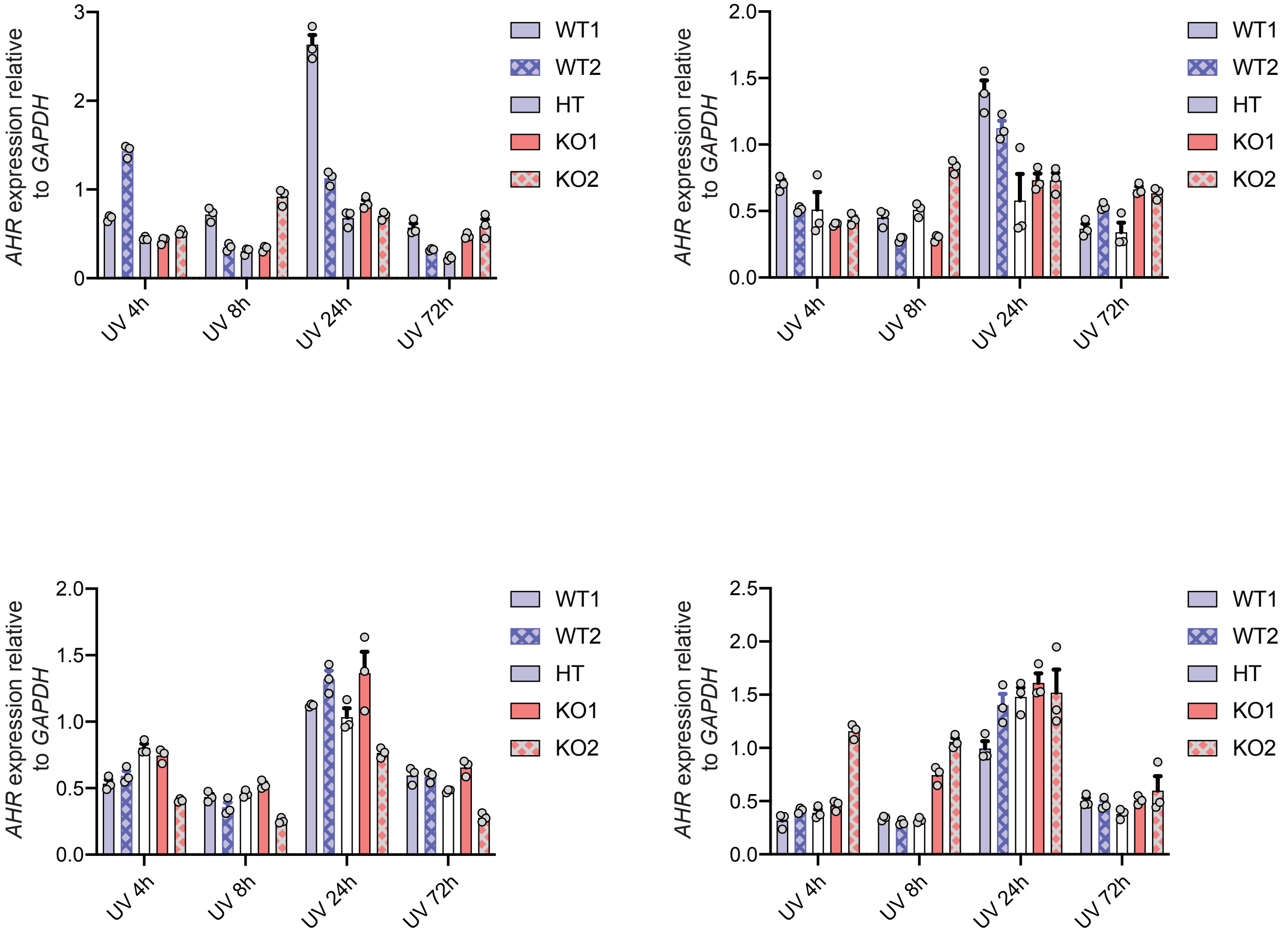

**Supplementary Figure 10: rs117132860-WT and rs117132860-KO cells show different cell proliferation under normal culture conditions.** (A) BrdU incorporation percentage was measured in five clones (2 WT, 2KO and 1 HT) growing under normal conditions, indicates a higher growth rate for WT cells and an intermediate rate for rs117132860-HT cells. Data shown are from two biological replicates. (B) Crystal violet quantitation of rs117132860-WT, rs117132860-KO, and rs117132860-HT clones grown for 1-4 days. One representative experiment with four biological replicates is shown, while three similar experiments were performed overall. The *P*-value is for the comparison of WT to KO clones (two-tailed paired Student's T-test) for the experiment shown; mean and SEM are graphed. The overall *P* value of 2 WT vs. 2 KO at D4 from 3 experiments (four replicates each) is 0.000019.

Supplementary Figure 10

A

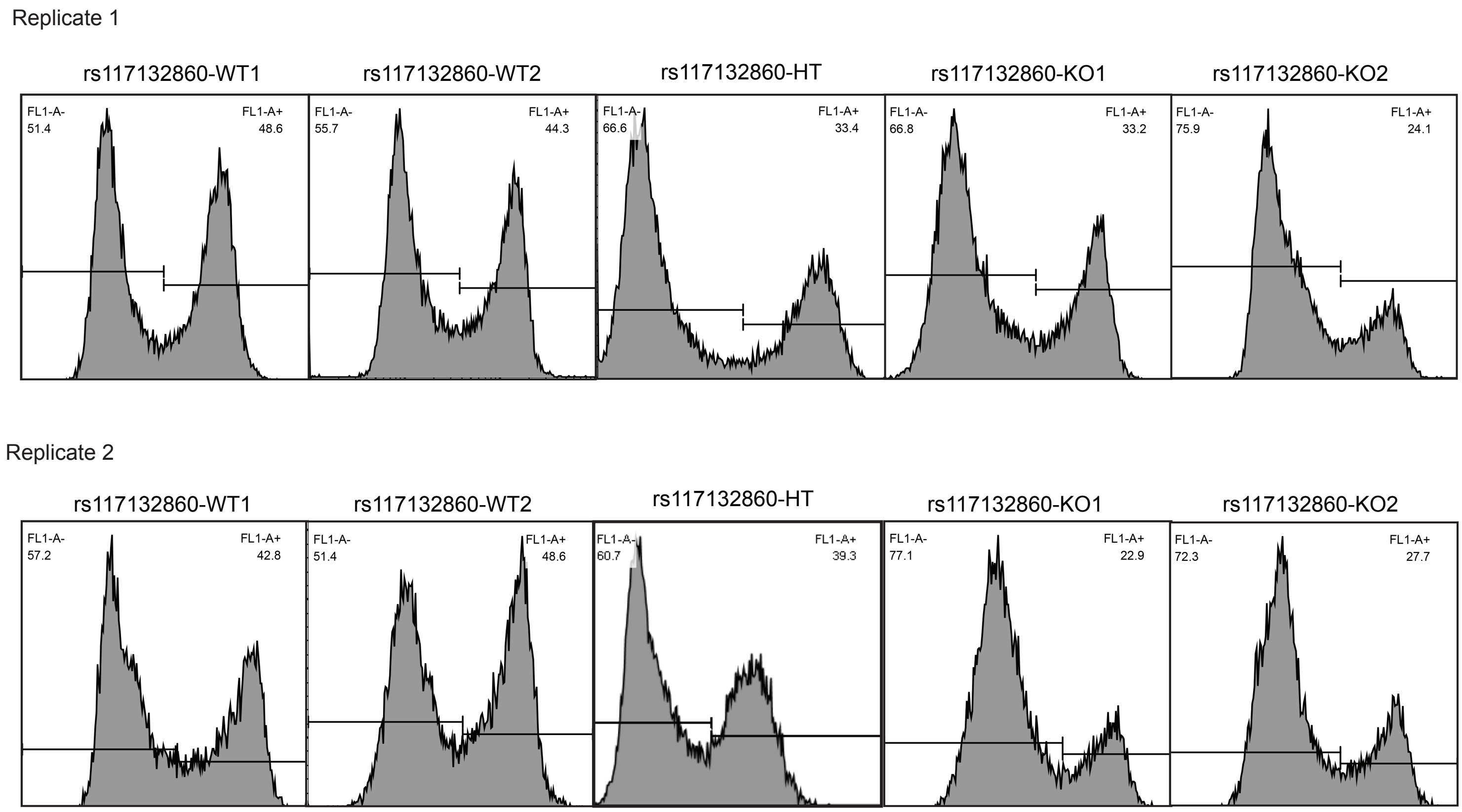


B

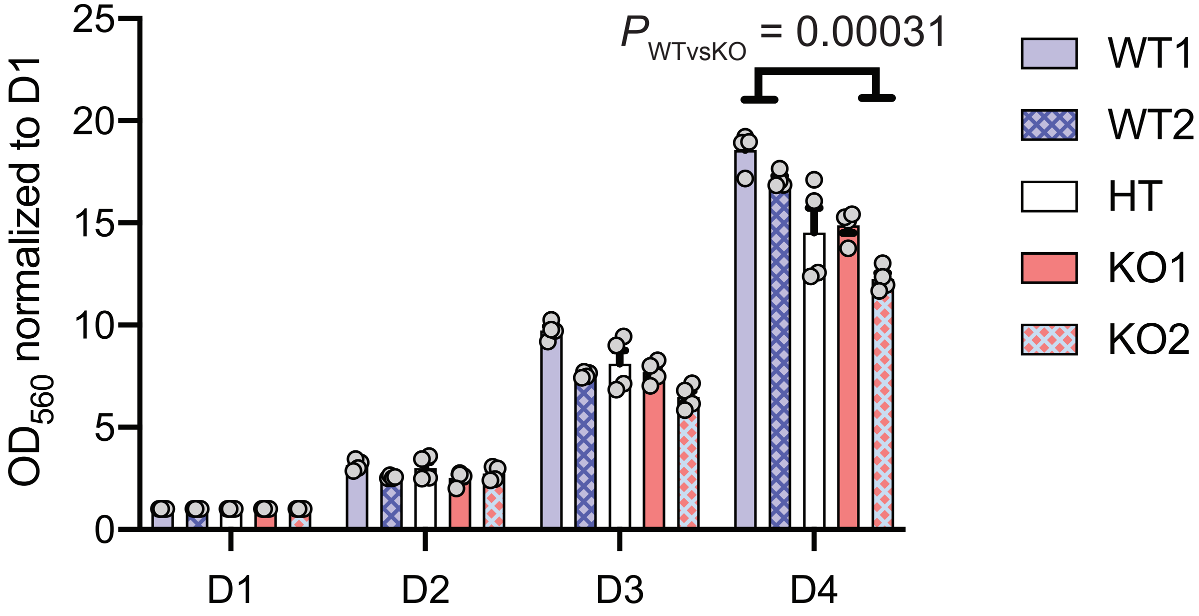

**Supplementary Figure 11: Crystal violet staining of additional CRISPR-Cas9 edited monoclonal melanocyte cell lines.** Crystal violet staining images of rs117132860-WT2, rs117132860-KO1, and rs117132860-HT cells not treated with UVB (24h), 72 hours after UVB treatment and Day 7 after UVB exposure followed by a zoomed-in image of day 7. The images shown here is from a representative experiment from three biological replicates.

### Supplementary Figure 11

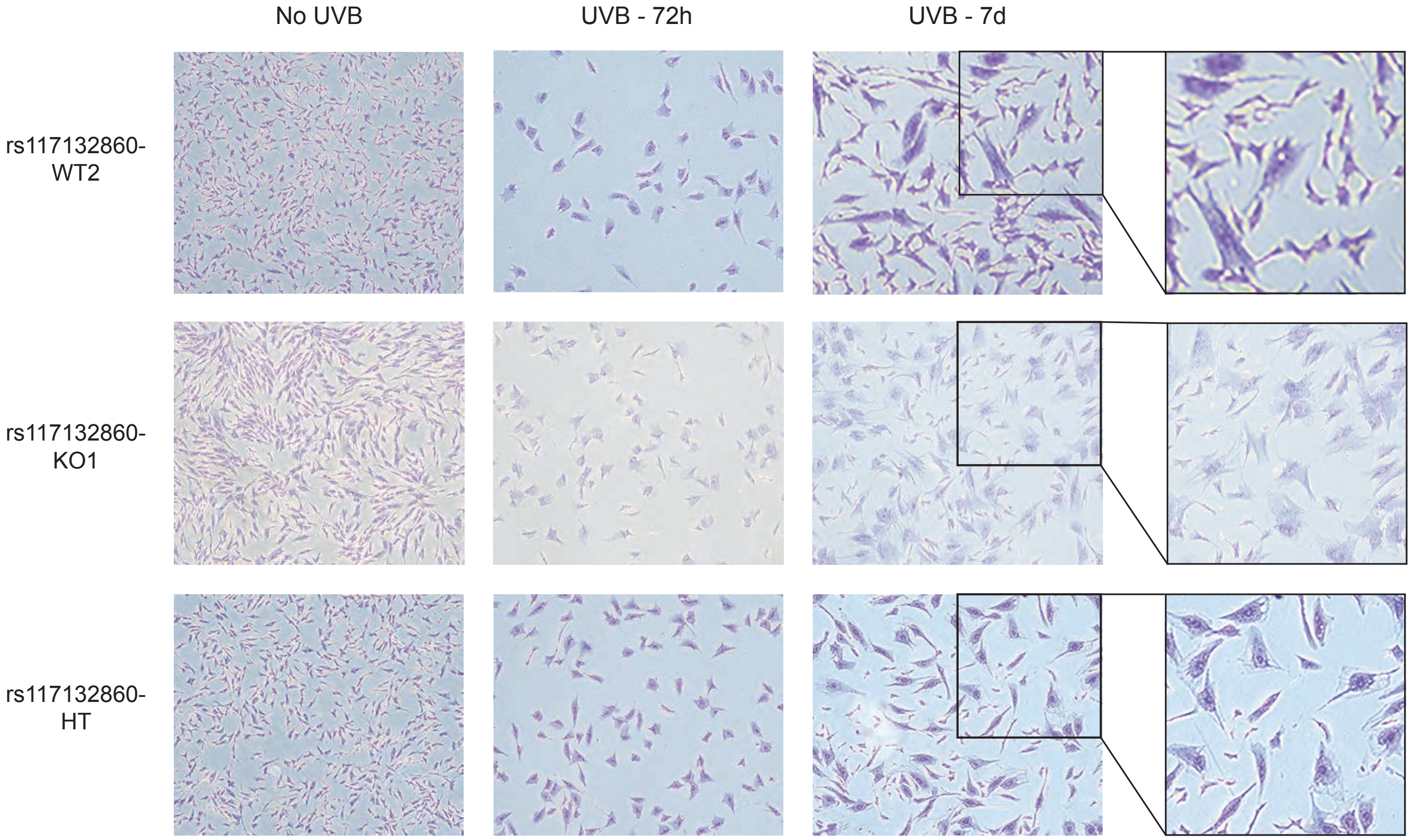

**Supplementary Figure 12: FACS analysis of additional CRISPR-Cas9 edited monoclonal melanocyte cell lines following UVB exposure.** FACS analysis of rs117132860-WT2 and rs117132860-KO1 clones, as well as rs117132860-HT cells, at day 7 after UVB treatment. Forward-scatter and side-scatter image indicates a mixed population with, BrdU and 7-AAD staining characterizing the cell cycle status of each population. The images shown here are from a representative experiment from three biological replicates.

### Supplementary Figure 12

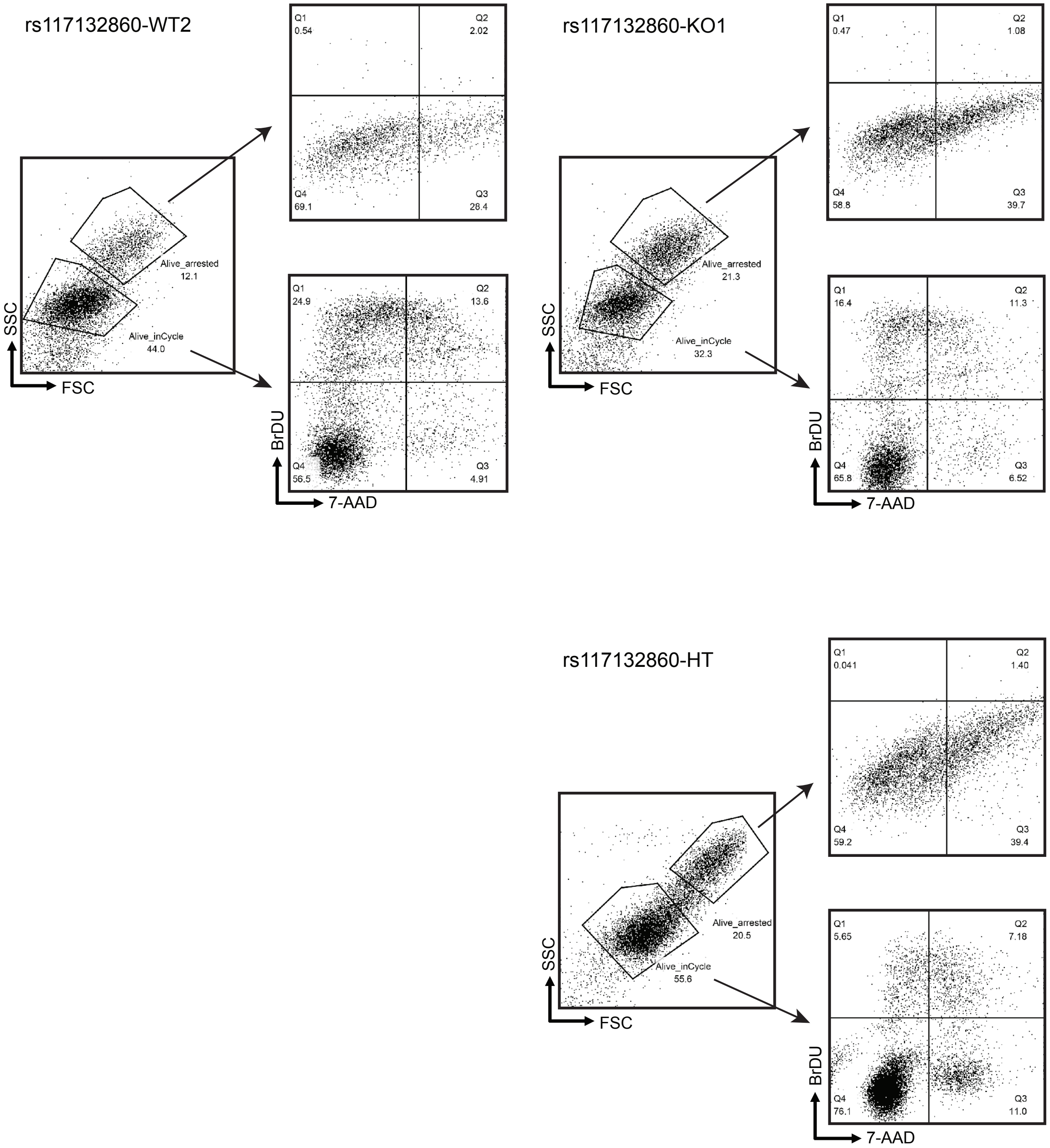
